# An ancestral pronephric contribution reveals the multilineage origin of the teleost gonad and revises the evolution of vertebrate gonadogenesis

**DOI:** 10.64898/2026.08.31.748213

**Authors:** Alexandra Depincé, Chloé Mayère, Francisca Hervas-Sotomayor, Aurélie Le Cam, Aurélien Brionne, Claudia Bevilacqua, Fany Blanc, Serge Nef, Manfred Schartl, Florent Murat, Amaury Herpin

## Abstract

Challenging the paradigm that pronephric field contribution to gonadal formation would be an amniote innovation, we demonstrate this trait is ancestral to bony vertebrates. Using cell lineage tracing, single-cell and spatial transcriptomics, and functional validation, we show that the teleost gonad arises from three distinct embryonic tissues, the pronephros, the coelomic epithelium, and the lateral plate mesoderm, in contrast to amniotes. This multi-tissue origin generates an unexpected lineage-based cellular diversity. Further cross-species comparisons over medaka, mouse, chicken and turtle unravel how lineage-specific deviations shape early gonadal development. Specifically, we map these variations amongst the different gene regulatory networks, outlining their physiological implications for specialized gonadal functions. Our results support a model in which heterochronic shifts are coupled to regulatory rewiring of conserved gene networks, driving lineage-specific developmental trajectories through a canalized developmental system drift.

## Introduction

Beneath their conserved architecture, comparative analyses of gonadal ontogenesis across vertebrates reveal an often under-appreciated complexity, driven by ancestral and derived traits, convergent evolutionary mechanisms, and dynamic species-specific trajectories of lineage specification.

In mammals, the gonadal primordia of both sexes emerge as pairs of thickened rows of coelomic epithelial cells ventral to the mesonephros. Following, bipotential gonads develop from a medulla/cortex structure (see [1] for review). The archetypal mammalian testis derives from embryonic testis cord structures made of germ and somatic supporting (Sertoli) cells, all surrounded by a layer of peritubular myoid cells. Concurrently, steroidogenic Leydig cells populate the interstitium between blood vessels and other connective tissues to give rise to the seminiferous tubules [1]. Analogously, typical mammalian ovaries develop a conserved characteristic cortical-medullary structure. The cortex, which includes granulosa, theca and stromal cells, is primarily derived from the proliferative coelomic epithelium (CE). In contrast, the medullary region originates from the sex cords interspersed with mesenchymal cells [1,2]. Interestingly, beyond these core processes, two additional mechanisms have been reported to further contribute to the cellular complexity and structural organization of the gonads: (***i***) ingression of multipotent cells from the CE [3–5], and (***ii***) migrating cells from the adjacent mesonephros [6–8].

From an evolutionary perspective, gonadal cord formation in mice proceeds through a distinct mechanism involving *de novo* aggregation of individual cells that ingress into the gonad after delamination of the CE and fragmentation of the basal lamina [8,9]. This is in contrast to turtles and anamniotes for which gonadal primordia are believed to develop after the sole invagination of the CE and preservation of the integrity of the basal lamina [9]. As a result, the contributions of either multipotent cells from the CE, or migrating mesonephric cells –thought to be key players in mammalian and avian gonadal development-have been largely overlooked if not denied outside mammals and birds [9] (see Figure 1A).

**Figure 1.**
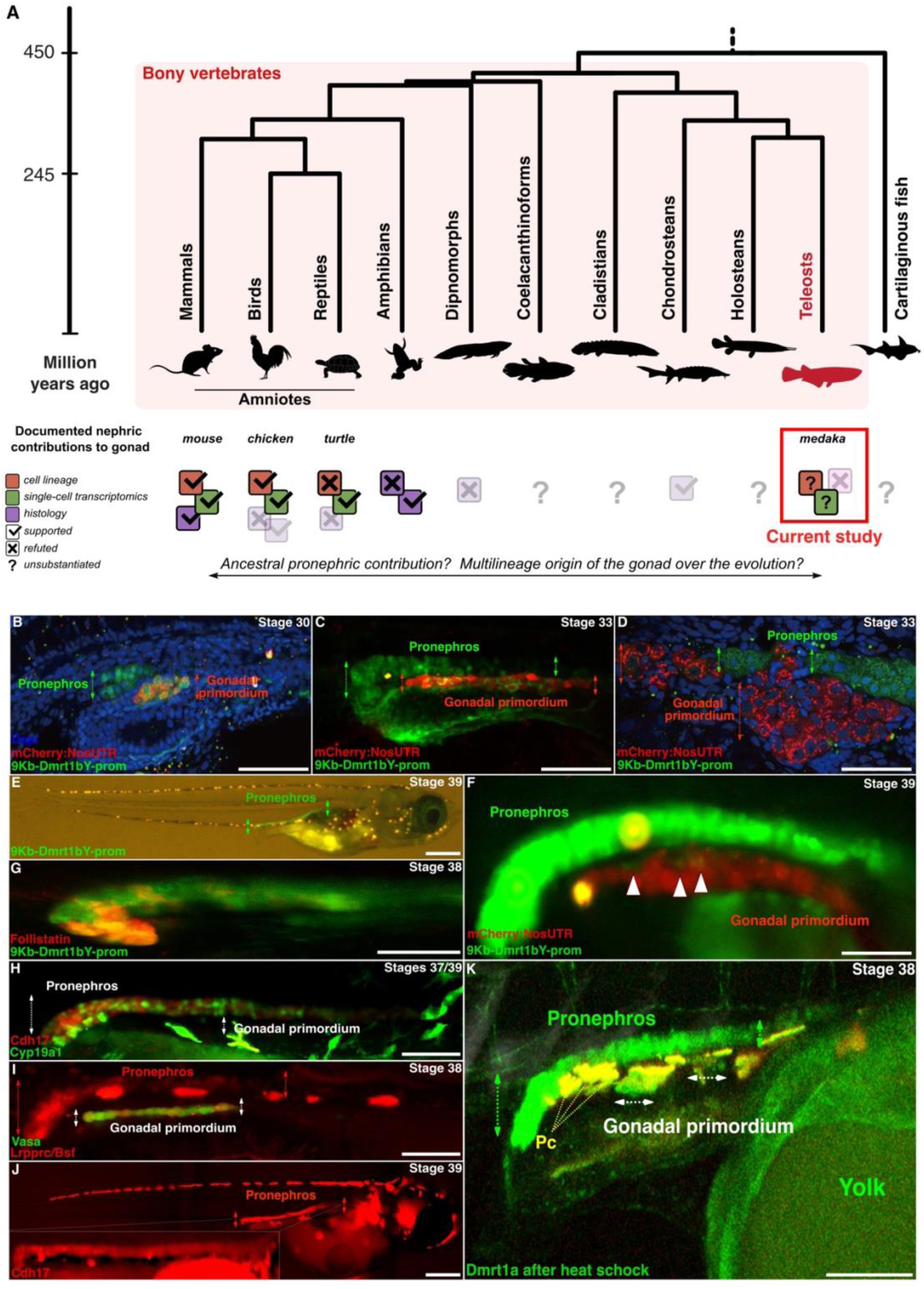
Phylogenetic relationship, pro-meso-meta-nephric contributions to gonads in vertebrates. **(A)** Current knowledge of pro- / meso-nephric contributions in bony vertebrates either supported, refuted or unsubstantiated by cell lineage tracing (brown square), single-cell transcriptomics (greenish square) or histological (purple square) analyses. Transparent items indicate studies for which definitive conclusions remain open. Documented pro/meso-nephric contributions to gonad: mice [7,64,69,71], chicken [50,55,72,73], turtle [9,10,57], amphibians [11–15,23–27,29], dipnomorphs [31], chondrosteans [32], medaka [21,22]. **(B to F)** GFP fluorescence driven by a 9-kb upstream *dmrt1bY* promoter is detectable in both the gonadal primordium and pronephros from stage 30 (3 days and 10 hours of development) through stage 39 (hatching, 8 days post-fertilization). *mCherry:NosUTR* fluorescence (red) specifically labels germ cells in the gonadal primordium (germ cells in red, **B, C, D and F**). Scale bars: **B and C**, 125 µM; **D**, 50 µM; **E**, 500 µM; **F**, 75 µM. **(G to I)** At hatching stage, *follistatin* (*fst* BAC reporter; **G**), gonadal *aromatase* (*cyp19a1* BAC reporter; **H**), and *bicoid stability factor* (*bsf/lrpprc*, promoter reporter; **i**) markers are co-expressed together with *cdh17* in the pronephros. Scale bar, **G**, 100 µM; **H**, 150 µM; **I**, 125 µM. **(J)** A 1.7 Kb upstream *cadherin-17* promoter drives mCherry expression specifically in the pronephros at hatching stage. Scale bar, 500 µM. **(K)** Heat shock of pre-hatching embryos induces ectopic synexpression of *dmrt1a* (BAC reporter) in both the gonadal primordium and pronephros at hatching stage. Pc, autofluorescent pigment cells. Scale bar, 200 µM.

Indeed, a large volume of classical literature has supported the paradigm that a mesonephric contribution to the gonads would be restricted to mammals and birds: (***i***) In turtles, either xenografts ([9]) or tissue cultures experiments ([10]), led to the conclusions that testis cord development occurs independently of the mesonephros. (***ii***) In amphibians, the origin of medullary cells from proliferating CE surrounding the gonad suggested that cells of the gonadal medulla are equivalent to Sertoli cells and follicular cells [11–15]. (***iii***) In fish the general understanding is that teleost gonads only originate from the peritoneal wall [16–20], with no evidence of a medulla derived from interrenal or mesonephric blastema. Specifically, in medaka, the gonadal primordium has been shown to develop simultaneously with the formation of the CE and cavity, making the contribution of the pronephric/mesonephric cells to the gonad “unlikely” [21,22].

However, challenging that general picture, several studies in anamniotes report, or suggest, a possible permeability between gonadal primordia and pro-/meso-nephros (Figure 1A): (***i***) In amphibians, where sex cords are absent, the source of medullary cells remains debated. While the origin of these cells from the proliferating CE is reported for canonical anuran models, a mesonephric blastema/tubules filiation is suggested in species representing more basal lineages [23–29]. More puzzling, in a caecilian, although late gonadal development takes places without the visible participation of the mesonephros, the initial gonad was reported to develop from rudimentary pronephric nephrostomial tubules [30]. (***ii***) Conversely, in lungfish, testicular tissues invade the kidney posteriorly [31]. (***iii***) Finally, histological studies in the sterlet proposed that the somatic cells of the gonad develop from the medial lips of nephrostomes, being clearly of opisthonephric (mesonephric) origin [32] (Figure 1A and Supplemental Figure 1). Collectively, these findings underscore the diverse and sometimes paradoxical contributions of nephrostomial structures to gonadal development across vertebrate lineages.

On a broader scale, while the general architecture of the adult testis and ovary is very similar across different phylogenetic groups, the underlying ontogenetic processes of morphogenesis and cell differentiation appear to be more plastic and obviously less conserved than initially thought [1,33]. Hence, being apparently a constant feature of the construction of the gonadal primordium phenotype, cell migration and ingression can be inferred to rely on ancestral and “evolutionary conserved” characteristics common to all vertebrates. Yet, from an evolutionary biology perspective, no clear picture emerges about ancestral versus derived features and mechanisms. Thus, no clear framework to reconcile lineage-specific deviations and adaptive inflexions exists.

To advance the understanding how gonadal ontogenesis evolved in vertebrates, we addressed the question of a possible pronephric contribution to the gonad in a teleost species. Using the medaka fish model, we performed *in vivo* fluorescent cell lineage tracing of early pronephric cells, monitored gonadal formation in the physiological situation and after pronephros development has been experimentally compromised, and conducted single-cell, and spatial transcriptomics to resolve the origins and fates of cell populations contributing to the medaka gonad. Extending this framework across species, molecular signatures of migrating pronephric cell populations were compared at single-cell resolution among mouse, chicken, turtle and fish. The comparative analysis enabled discerning ancestral *versus* derived features, and allowed reconstructing the conserved and divergent evolutionary trajectories underlying early gonadal developmental processes in vertebrates.

## Results

### Expression patterns of gonadal and pronephros markers suggest pronephric contribution to gonad development in medaka

The gonad of medaka has been reported to develop through the coordinated differentiation of two distinct cell lineages: germ cells and the surrounding somatic gonadal mesoderm [22]. Once specified, primordial germ cells (PGCs) remain closely associated with endodermal tissues and migrate *via* the dorsal gut mesentery to the region of the presumptive gonad [21], which is derived from the genital ridge epithelium. However, unlike in mammals, no indication of a medullary tissue could be identified in the teleost gonad. Prior to differentiation, all somatic cells appear to be derived from a cortical epithelial layer and are morphologically similar in males and females [16–20]. Shortly before hatching, when *dmrt1bY*, the male-determining gene in medaka, is expressed in the male gonad primordium, the germ cells in the female gonad actively proliferate and undergo meiosis, while this occurs much later in male gonads [34].

For studying early medaka gonadal differentiation and ontogenesis, we established several transgenic fluorescent reporter lines to follow the dynamics of expression of several gonadal markers at cellular resolution during gonadal commitment (Figure 1B to 1K). Resulting in optimal spatial resolution together with high reliability of gene expression [35,36], these fluorescent lines can be expected to recapitulate all different cell lineages constitutive of the primordial gonads: (***i***) pre-supporting, i.e. pre-Sertoli or pre-granulosa (*dmrt1bY*, *dmrt1a*, and *follistatin*; Figure 1B to 1G)-, (***ii***) steroidogenic (*cyp19a1*; Figure 1H) and (***iii***) germ cell (*bsf/lrpprc*; Figure 1I) lineages. A reference line expressing Cadherin-17 promoter-driven mCherry fluorescence specifically in the pronephros was generated (Figure 1J)

As early as stage 30 (3 days and 10 hours of development, [37]) and up to hatching stage (stage 39), *dmrt1bY* is not only expressed in the pre-supporting cells of the gonadal primordium in between the germ cells [34,38], but also in the forming pronephros (Figure 1B to 1F).

Concomitantly, the *cyp19a1 aromatase* reporter fluorescence – a marker of the steroidogenic lineage - is detected in a homogenously distributed subset of cells all along the pronephric tube (Figure 1H compared to 1J). At same stages, *follistatin*-reporter expression localises to a restricted number of cells at the very posterior tip of the pronephros (Figure 1G). Finally, *Bsf*/*Lrpprc*, a germ line marker [39], is, in addition to a subset of germ cells, also expressed in cells belonging to the most posterior part of the pronephros (Figure 1I). Interestingly, heat shock in medaka embryos induces XX female-to-male sex reversal, marked by premature activation of *dmrt1a* in supporting cells of the primordial gonad and its ectopic expression in the posterior end of the pronephric duct (Figure 1K and [40]). That heat-induced synexpression suggests a functional connection between these tissues.

### *In vivo* cell lineage tracing reveals pronephric field contributions to both female and male gonads in medaka

To further address the question of a potential pronephric contribution to the gonads in medaka, we made use of the “brainbow” system [41] to perform cell lineage specific tracing of pronephric precursor cells *in vivo* (Figure 2A).

**Figure 2.**
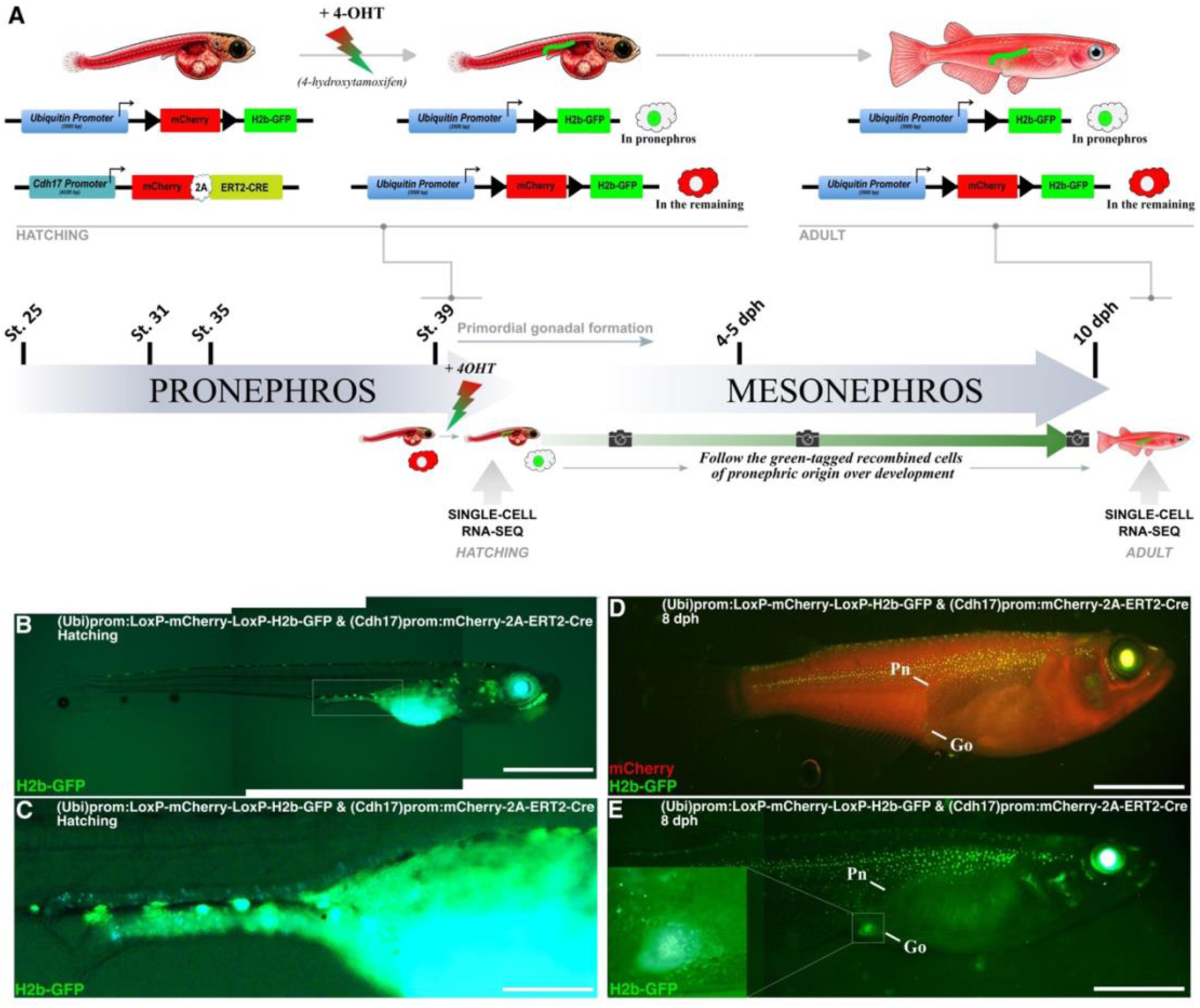
Generation of a *cdh17* medaka model for inducible *in vivo* cell lineage tracing of early pronephric field cells. **(A)** The *cdh17* medaka model carries two transgenes enabling pronephric cell-specific labelling: a switch from ubiquitous cytoplasmic-localized mCherry to pronephric-restricted nuclear-localized GFP fluorescence (red to green, respectively). Upon 4-OHT exposure, CRE recombinase - expressed under the pronephric-specific *cdh17* promoter - translocates to the nucleus, inducing recombination and excision of the mCherry-stop cassette. This results in permanent GFP labelling of pronephric cells. In absence of 4-OHT, CRE recombinase remains cytoplasmic, and no recombination (GFP fluorescence) is detected (see Supplemental Figure 2). Following pronephric-restricted recombination induction shortly before hatching, GFP fluorescence was monitored (cameras) at different developmental stages: hatching stage **(B and C)**, 8 dph **(D and E)** and adulthood (see Figure 3). Scale bars, **B** and **C**, 450 µM; **D and E**, 750 µM.

Briefly, we first established a “reporter” line (*Ubi prom:LoxP-mCherry-LoxP:H2B-GFP*) that is able to switch from a ubiquitous cytoplasmic mCherry (red fluorescence) to a specific nuclear-localized GFP (green fluorescence) expression after controlled homologous recombination. In this line, a 3.5 kb-long ubiquitin promoter drives the ubiquitous expression of a mCherry-stop cassette flanked by *LoxP* sites. This cassette is followed by the open reading frame coding for the nuclear-addressed GFP (H2B-GFP). A second “driver” line (*Cdh17prom:mCherry-2A-ERT2-Cre*) expresses the ERT2-Cre ((4-OHT), tamoxifen-inducible recombinase) fused to a 2A self-cleaving system [42] with mCherry under the control of the Cdh17 pronephros-specific promoter (Figure 2A, see also Figure 1J for Cdh17 promoter specificity). After breeding of these two lines (*Ubi prom:LoxP-mCherry-LoxP:H2B-GFP/ Cdh17prom:mCherry-2A-ERT2-Cre*), incubation of hatchlings with 4-OHT will provoke the cytoplasmic-to-nucleus shuttling of Cre and specific recombination in the pronephros. The mCherry-stop cassette is then specifically excised and H2B-GFP expression is only occurring in the pronephros and hence cell tracing is activated (Figure 2A to 2C and Supplemental Figure 2A to 2C). In absence of 4-OHT induction, no illegitimate recombination (GFP fluorescence outside of the pronephros) was detected (Supplemental Figure 2D to 2F). This approach enables real-time tracking of pronephric precursor cell migration and fate commitment throughout development (Figure 2B to 2E). We observed that by 8 dph, GFP-positive cells that underwent recombination in the pronephros at hatching stage migrate to the gonad. The significantly stronger gonadal fluorescence, compared to the pronephros, suggests active proliferation of these cells within the gonad (Figure 2D and 2E). Still at adult stage, cells of pronephric origin (recombined cells expressing H2B-GFP) were present in both female (Figure 3A to 3D) and male (Figure 3G to 3L) gonads.

**Figure 3.**
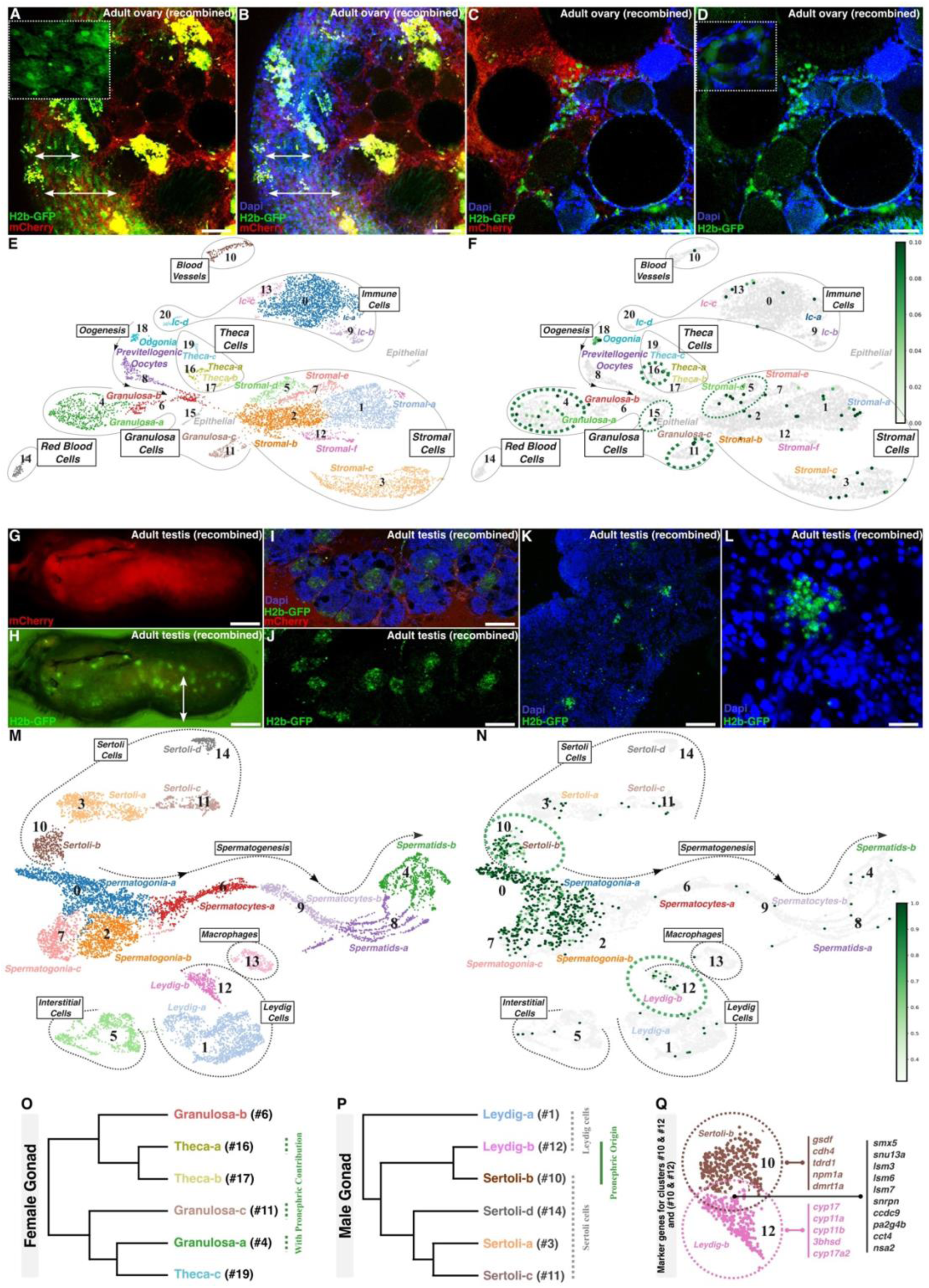
*In vivo* cell lineage tracing and single-cell transcriptomic analyses reveal contribution of the pronephric field to both female and male adult medaka gonads. **(A to D and G to L)** Medaka embryos subjected to pronephric-specific recombination around hatching stage display GFP-positive cells (nuclear-localized fluorescence) of pronephric origin in both female **(A to D)** and male **(G to L)** adult gonads. **(A to D)** <u>Ovary</u>: pronephros-derived cells (green fluorescence, arrows) contribute to the *tunica albuginea* (A and B), and supporting cell populations around young oocytes, including granulosa, theca and stromal cell types **(C and D)**. **(G to L)** <u>Testis</u>: pronephros-derived cells (green fluorescence) integrate into supporting cell populations within germinal cysts **(I to L)**. (**E and F**, and **M and N**) Single-cell transcriptomic analyses and UMAP projections of adult female **(E and F)** and male **(M and N)** gonads, following pronephric lineage tracing at hatching stage, elucidate the molecular fate of pronephric-contributing cells. The color bar indicates GFP-expression strength. **(E and F)** <u>Ovary</u>: GFP mRNA-expressing cells of pronephric origin **(F)** contribute to the epithelial and stromal layers, as well as sub-populations of granulosa and theca cells **(F)**. **(M and N)** <u>Testis</u>: cells of pronephric origin (expressing the GFP mRNAs; **(N)**) contribute to sub-populations of Sertoli and Leydig cells **(N). (O and P)** Hierarchical clustering dendrograms of female **(O)** and male **(P)** gonadal cell sub-populations. **(Q)** Top-Differentially Expressed Genes (DEG) shared and specific to Sertoli and Leydig sub-populations of pronephric origin.

In the adult female gonad, cells of a pronephric origin are located around the ovary just internal to the most external germinal epithelium layer. This layer corresponds to the *tunica albuginea* (arrows in Figure 3A and 3B). Within the deeper regions of the ovary, H2B-GFP-expressing cells are observed surrounding small oocytes in the germinal cradles (Figure 3C and 3D). These cells represent newly differentiated supporting cells of the granulosa lineage. The observation that nearly all cells surrounding specific oocytes express GFP fluorescence strongly suggests a clonal origin of that supporting cell population (insert in Figure 3D).

Further investigation using single-cell analysis of adult ovaries from hatchlings that underwent recombination (Figure 3E and 3F; Supplemental Figure 3A and 3B) revealed that, among the three-identified granulosa and theca cell types, specific sub-types (granulosa-a, cluster #4; granulosa-c, cluster #11; and theca-a, cluster #16, all expressing GFP mRNAs) are of pronephric origin. Consistent with our *in-situ* observations (Figure 3A to 3D), pronephric field contribution is also apparent for the epithelial (cluster #15) and stromal (stromal-d, cluster #5) layers. Interestingly, hierarchical clustering dendrogram of these granulosa and theca sub-populations reveals no lineage-based correlation among cells of pronephric origin (Figure 3O). Instead, the apparent fate-base clustering supports the interpretation that cells from different origins all contribute to the different sub-populations, regardless of their developmental source (Figure 3O).

In the adult male gonad, cells of pronephric origin are distributed in the inner part of the testis (Figure 3G and 3L). H2B-GFP-expressing cells contribute to the interstitial tubules (Figure 3I to 3K), as well as to the supporting cell lineage of the germinal lobules (Figure 3I to 3L). The observation that most supporting cells within a given germinal lobule express GFP indicates that these cells are likely clonal (Figure 3L). Furthermore, single-cell analysis of adult testes from hatchlings that underwent recombination enabled the characterization of six distinct subtypes of supporting cells: four Sertoli (Sertoli-a, cluster #3; Sertoli-b, cluster #10; Sertoli-c, cluster #11; Sertoli-d, cluster #14) and two Leydig subtypes (Leydig-a, cluster #1; Leydig-b, cluster #12; Figure 3M and 3N; Supplemental Figure 3C and 3D). Notably, only the Sertoli-b and Leydig-b populations are of pronephric origin (Figure 3N). While early spermatogonia populations do express GFP mRNA - suggestive of a pronephric origin -, this expression is lost in differentiating spermatocytes and then spermatids (Figure 3N). This pattern most likely indicates RNA transfer from the Sertoli-b lineage of pronephric origin, rather than genomic recombination in the germinal lineage and pronephric origin. In contrast to the ovary, hierarchical clustering reveals a robust lineage-based correlation among cells of pronephric origin (Sertoli-b and Leydig-b; Figure 3P) in testis. This suggests that the subpopulations of Sertoli and Leydig cells derived from the pronephros, do not only supply the general pool of supporting cells (as observed in the ovary), but also retain a molecular memory of their embryonic origin, expressing a similar set of genes mainly related to RNA processing and splicing (Figure 3Q). Spatial prediction of single cell clusters corresponding to these different subpopulations further demonstrate distinct and specific testicular localizations (dispersed: Sertoli-a; peripheral: Sertoli-b; peripheral/dispersed: Leydig-b; or more central: Sertoli-c and –d) according to their subtypes (Figure 4A).

**Figure 4.**
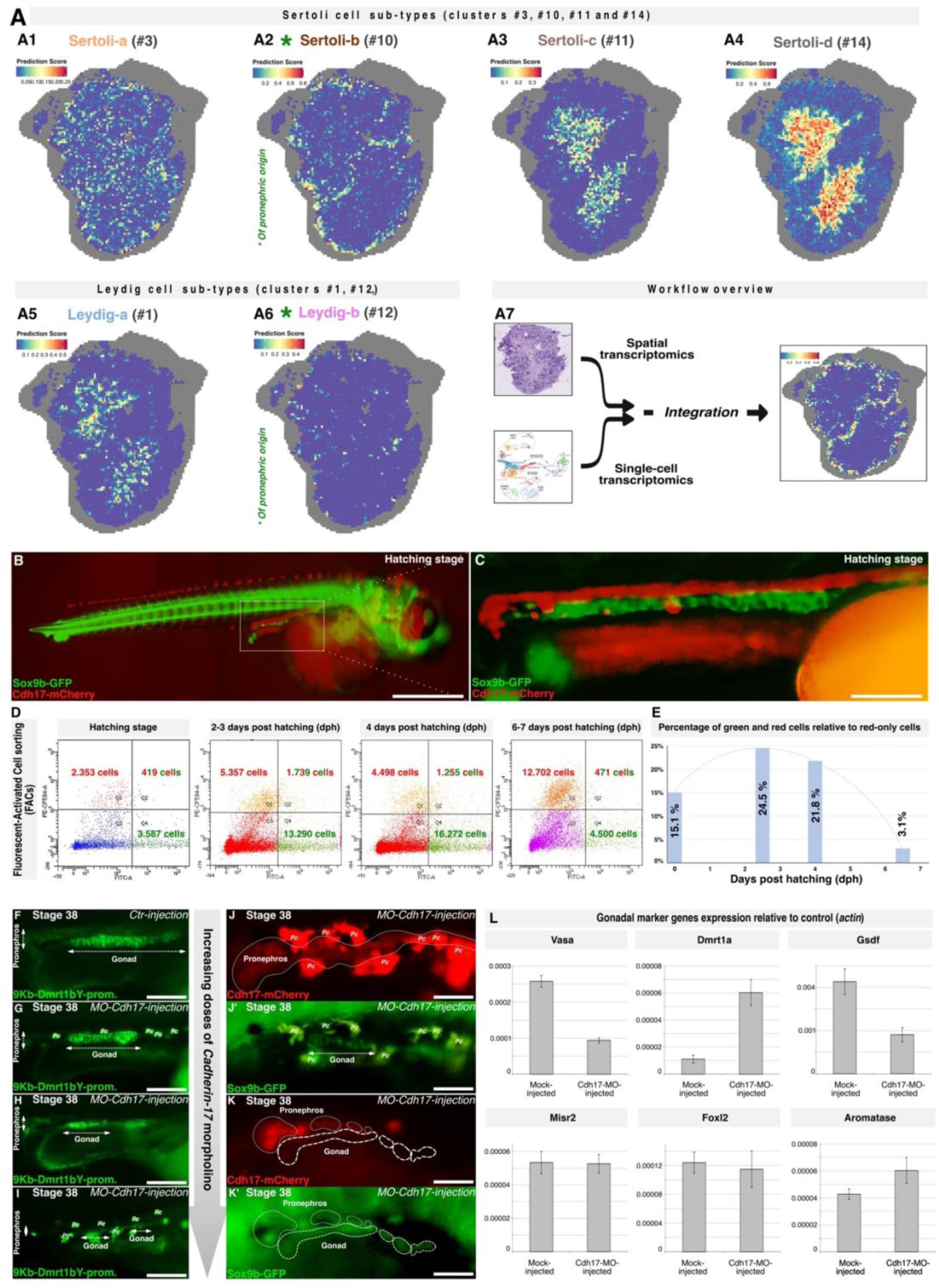
Spatial testis transcriptomic analyses and kinetics of nephric contribution to gonad formation under impaired pronephros development. **(A)** Scheme of the spatial transcriptomics experiment (Visium HD 10x) showing the level of the medaka testis cross section. **(A1 to A6)** Visium 10x slide representations showing spatial predictions of single-cell clusters corresponding to Sertoli **(A1 to A4)** and Leydig **(A5 and A6)** sub-clusters (clusters #3, 10, 11, 14 and 1, 12 respectively) as defined in Figure 3E and 3M. Each dot represents a 16 µm capture area. **(A7)** workflow’s overview. **(B to E)** Medaka embryos expressing red (pronephros) and green (gonadal primordium) fluorescence were dissected for the nephro-gonadal region **(B and C)**, cells dissociated and then subjected to FACS sorting for retrieving red, green and red/green populations **(D)**. Over early development, the red/green cell population peaks shortly after hatching and persists up to 2-3 dph **(E)**. **(F to L)** Impairing pronephros development hinders normal gonadal development at hatching. **(F)** Control morpholino injection. **(G to I)** Increasing doses of *cdh17-morpholino* (*cdh17-mo*) progressively inhibit gonadal development as visualized by *dmrt1bY*-reporter fluorescence. **(J to K’)** Gradual impairment of cdh17-mo-induced pronephros development proportionally affects early gonadal formation. **(L)** Expression of early gonadal primordium markers is differentially altered following impaired pronephric formation. Scale bars, **B**, 750 µM; **C**, 150 µM; **F** to **I**, 120 µM; **J to K’**, 60 µM.

### In medaka, the pronephric field’s contribution to the primordial gonad peaks at 2-3 days post hatching

Depending on species, vertebrates display up to three kidney component categories relative to development and function [43]: (***i***) the pronephros, (***ii***) the mesonephros (referred as opisthonephros when acting as a functional kidney), and (***iii***) the metanephros (being the “functional” kidney in amniotes). Anamniotes and other *Sarcopterygii*, *Actinopterygii* (including teleosts) and cartilaginous fishes, pass through the first two types at different stages of ontogenesis (see Supplemental Figure 1 and [44,45] for review). While reptiles possess a metanephric kidney as the terminal renal organ, they uniquely maintain kidney progenitor cell populations throughout life and continually develop new nephrons [46]. In mammals and birds, only the mesonephric contribution is documented (see [1] for review, and [47]). As early gonadal primordium commitment and development span over pro-to-meso-nephric transition in medaka (from hatching to 8 dph), we explored the temporal dynamics of this nephric migration. A double transgenic reporter line, specifically expressing (***i***) the green fluorescence (GFP) in the somatic part of the primordium gonad due to the sox9b promoter [48], and (***ii***) the red fluorescence (mCherry) in the pro/meso-nephros under control of the Cadherin-17 promoter was generated (Figure 4B and 4C). Gonadal and pro/meso-nephros areas of 300 hatchlings at specific developmental stages of development were micro-dissected, subjected to cell dissociation and finally engaged to Fluorescent-Activated Cell Sorting (FACS; (Figure 4D). Counting the different fluorescent cell populations (green-only, red-only and green/red) over the different time points (hatching, 2-3, 4 and 6-7 dph) revealed a higher proportion of double fluorescent (red and green, 24.5%) cells 2-3 dph while a fully functional pronephros is operative (Figure 4D and 4E). Later on, when the pronephros is resorbed and the mesonephros develops (4 to 5 dph onward) the proportion of the double fluorescent cell population steadily decreases from 24.5% 2-3 dph down to 3.1% at 6-7 dph (Figure 4E). In summary, the primary nephric contribution to the gonads in medaka appears to occur between hatching and 2-3 dph stages, corresponding to the pronephric ontogenic developmental form of the kidney (opisthonephros; Supplemental Figure 1). The presence of only 3.1% double-positive fluorescent cells at 6-7 dph, likely reflects fluorescence persistence rather than new recruitment. Consequently, a significant later contribution from the mesonephros can be largely excluded.

### Pronephric field contribution is essential for proper development of the primordial gonad in medaka

Whereas mesonephric cell migration has been shown to be necessary for male gonadal development in mice [7,49], such a contribution appears to be dispensable for proper gonadogenesis in birds [50,51] (see Figure 1A).

To functionally evaluate the requirement of the pronephric contribution for gonad development in medaka, we interfered with pronephros development (Figure 4F to 4K’). While constitutive knock-out of *cadherin-17* gene, a gene shown to be required to maintain pronephric duct integrity [52], is early lethal (CRISPR-Cas9, personal data), for that experiment, expression of *cadherin-17*, has been progressively knockdown after injection of increasing doses of a *Cdh17-morpholino*. Gonad formation was monitored using different fluorescent markers (Figure 4F to 4K’) and regulation of classical gonadal marker genes (*dmrt1bY*, *dmrt1a*, *foxl2*, *misr2*, *gsdf*, *cyp19a1* and *vasa*) was assessed by real time quantitative PCR (Figure 4L). Increasing doses of Cdh17-Mo progressively impaired gonad formation (Figure 4F to 4I), supporting a causal relationship between pronephric disruption and defective gonad development (Figure 4J and 4J’ compared to 4K and 4K’). Consistently, pronephric impairment differentially affected gonadal marker expression: whereas *dmrt1a*, *cyp19a1* and *vasa* were upregulated, *gsdf* was downregulated, and the expression levels of *dmrt1bY*, *foxl2* and *misr2* remained unchanged (Figure 4L). These results suggest that pronephric-derived cells selectively contribute to specific cell sub-populations.

### Single-cell transcriptomics reveals the molecular trajectories underlying cell fate transitions

To resolve the molecular dynamics of pronephric cells contributing to the gonad, we performed single-cell RNA-seq on hatching-stage medaka pronephros, gonadal primordium and pronephric-derived gonadal cells (Figure 5). Given the particularly low occurrence of the pronephric-contributing-to-gonad cell populations (see Figure 4D), sorted gonadal and pronephros-enriched cells for sox9b-GFP and cdh17-mCherry fluorescences respectively-were subjected to single cell RNA sequencing. Clustering analysis and UMAP visualization revealed 40 clusters encompassing the whole pronephros from proximal to distal tubules and further caudal to the cloaca (Figure 5A). Pre-supporting gonadal, male (Sertoli) and female (granulosa) gonadal as well as germ cell populations were also identified (Figure 5A; see Supplemental Figure 4A for the complete clustering and annotation). Cells of the coelomic epithelium (CE) and the lateral plate mesoderm (LPM) were also captured (Figure 5A and 5B). Few clusters remained unannotated (Figure 5A and 5B). Hierarchical clustering (Figure 5B) established a clear relationship between different sub-populations of gonadal/pre-gonadal supporting cell clusters (pre-granulosa/granulosa clusters #17/#8 and gonadal pre-supporting #14, granulosa #3/#5 and Sertoli #4) together with either coelomic epithelium (clusters #7/#36) or lateral plate mesoderm (clusters #9/#20), suggesting two additional distinct embryonic origins for the gonadal supporting cell lineage besides pronephros (Figure 5B). Interestingly, cells belonging to the two LPM clusters (clusters #9/#20) additionally express the *nr5a1*/*sf-1* nuclear receptor implicated in steroidogenesis (see [53] for review). With that respect, our analysis does not pinpoint clearly defined embryonic steroidogenic cell precursors *per se* (such as “fetal” pre-Leydig or pre-Theca cells) which have been classically characterized in mouse [54] and chicken [2,55]. However, key steroidogenic enzymes – including *cyp17*/c*yp17a2* and *star*-are highly expressed in clusters #4/#19 and #35 respectively (Figure 5C to 5F). While clusters #4/#19 display a clear Sertoli signature, the identity of cluster #35 remains undetermined. Such distribution of the “embryonic” steroidogenic function(s) among several cell types suggests that (***i***) analogous functions can be endorsed by different cell lineages, and (***ii***) part of the adult steroidogenic cells likely originate from embryonic supporting precursors, presumably derived from *nr5a1*/*sf-1*-expressing LPM cells.

**Figure 5.**
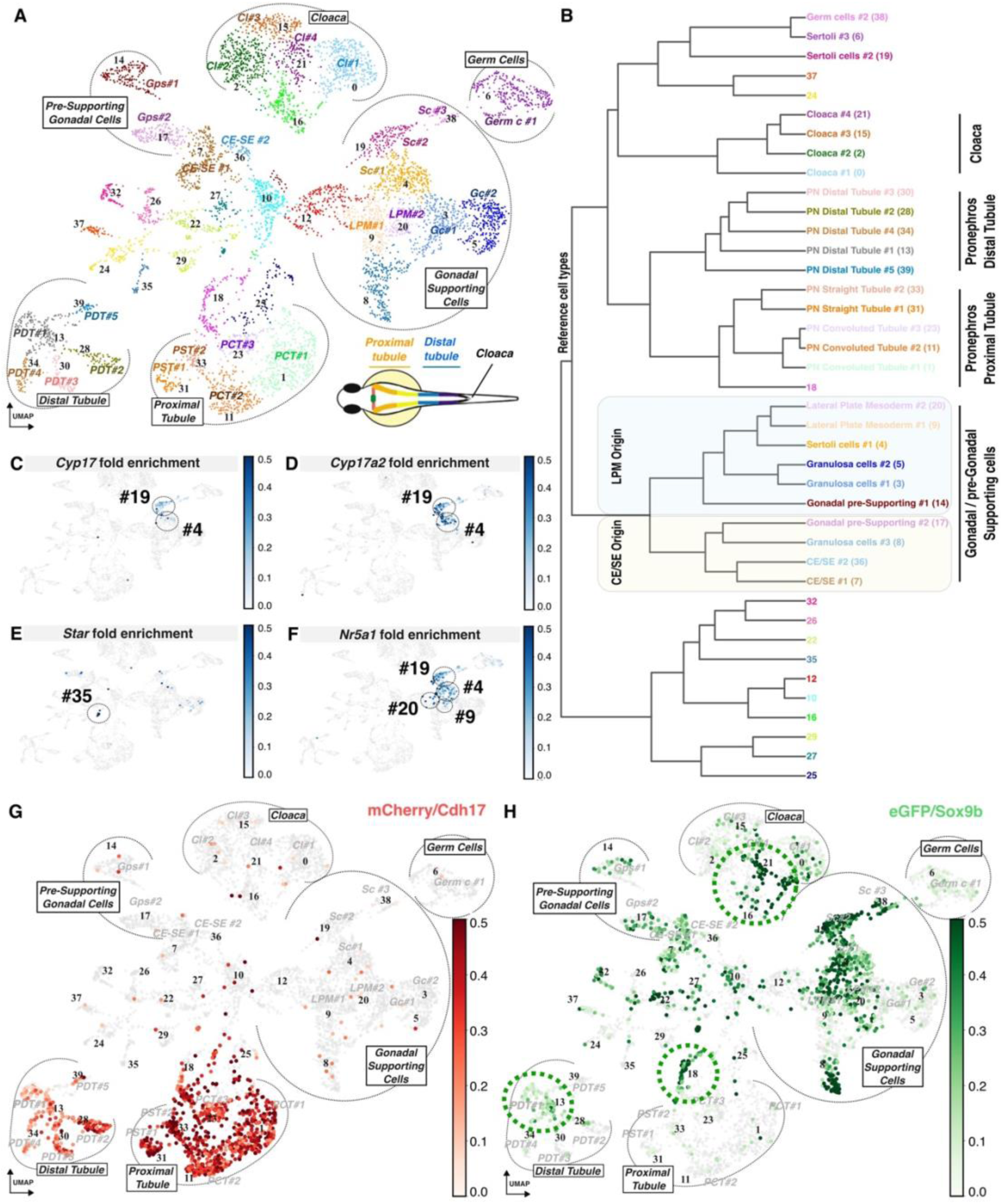
Single cell transcriptomic analysis of medaka pronephric, gonadal primordium and pronephric-contributing gonadal cells at hatching stage. **(A to I)** Mixed male and female hatching-stage primordial gonads and pronephros enriched for Sox9b-GFP and cdh17-mCherry–positive cells, respectively (see also Figure 4B to 4D and STAR methods) were subjected to single cell RNA sequencing. Clustering and Umap visualization of the 7,317 transcriptomes revealed 40 clusters encompassing the whole pronephros from proximal tubule (clusters #1/11/23/31/33) to distal tubule (cluster #13/28/30/34/39) and caudal to cloaca (clusters #0/2/15/21), pre-gonadal supporting cells (clusters #7/9/14/17/20/36), male and female gonadal (clusters #4/19 and 3/5 respectively), and gem cells (clusters #6/38) **(A and B)**. **(C to F)** Expression of early steroidogenic markers such as *cyp17*, *cyp17a2* and *Star* is restricted to clusters #4, 19 and 35. **(G and H)** The mCherry/Cdh17 markers are specifically expressed in the proximal and distal parts of the pronephros **(G)**. The GFP/Sox9b markers are expressed in the gonadal pre-supporting and supporting cells as well as in some pronephric cells (proximal and distal tubules, and cloaca) and other non-annotated clusters (clusters #16/18/22) **(H)**. Green dotted circles (clusters #13/16/18/21) highlight specific clusters co-expressing both pronephric mCherry/Cdh17 and gonadal eGFP/Sox9b signatures **(H and G compared to H)**. Color bars indicate fold enrichments **(C to H)**. LPM: Lateral Plate Mesoderm; CE-SE: Coelomic/Supporting Epithelium; Gc: Granulosa cells; Sc: Sertoli cells; Gps: Gonadal pre-supporting cells; Germ c: Germ cells; PDT: Pronephros distal Tubule; PST: Pronephros Straight Tubule; PCT: Pronephros Convoluted Tubule; CI: Cloaca.

Several unannotated clusters, despite lacking a definitive overall gonadal signature, do express the gonadal markers *sox9b*/*GFP*, potentially indicating a pre-gonadal fate state (Figure 5G and 5H). Particularly, few clusters express both the *cdh17* pronephric and the *sox9b* gonadal markers (see clusters #13 and #18 in Figure 5G and 5H), and clusters of cells belonging to the cloaca clearly express the sox9b early gonadal marker (see clusters #16 and #21 in Figure 5H). More intermediate clusters express both cdh17 and sox9 markers (see cluster #10, #12, and #22 in Figure 5G and 5H, see also Supplemental Figure 4B). Such cells or groups of cells expressing both markers actually confirm-at a molecular scale-our previous observations reporting the existence of cell populations expressing pronephros and gonadal markers (see Figure 1B to 1K). Hence, the scRNA-seq results further support the existence of multiple pronephric cell contributions in a stepwise manner early during gonadal ontogenesis in medaka (Figures 2, 3 and 4). This emphasizes on a larger scale the plasticity and complexity of cell fate transitions during gonadal development in vertebrates. Un-annotated clusters (Figure 5A and 5B), though expressing a slightly narrower transcript range than other clusters, lack identifiable molecular signatures (Supplemental Figure 4A). Notably, all these clusters - without any exception-are enriched for either motility or stemness signatures (or both, Figure 6 and Supplemental Figure 6), likely representing migrating cells in transition between fates.

**Figure 6.**
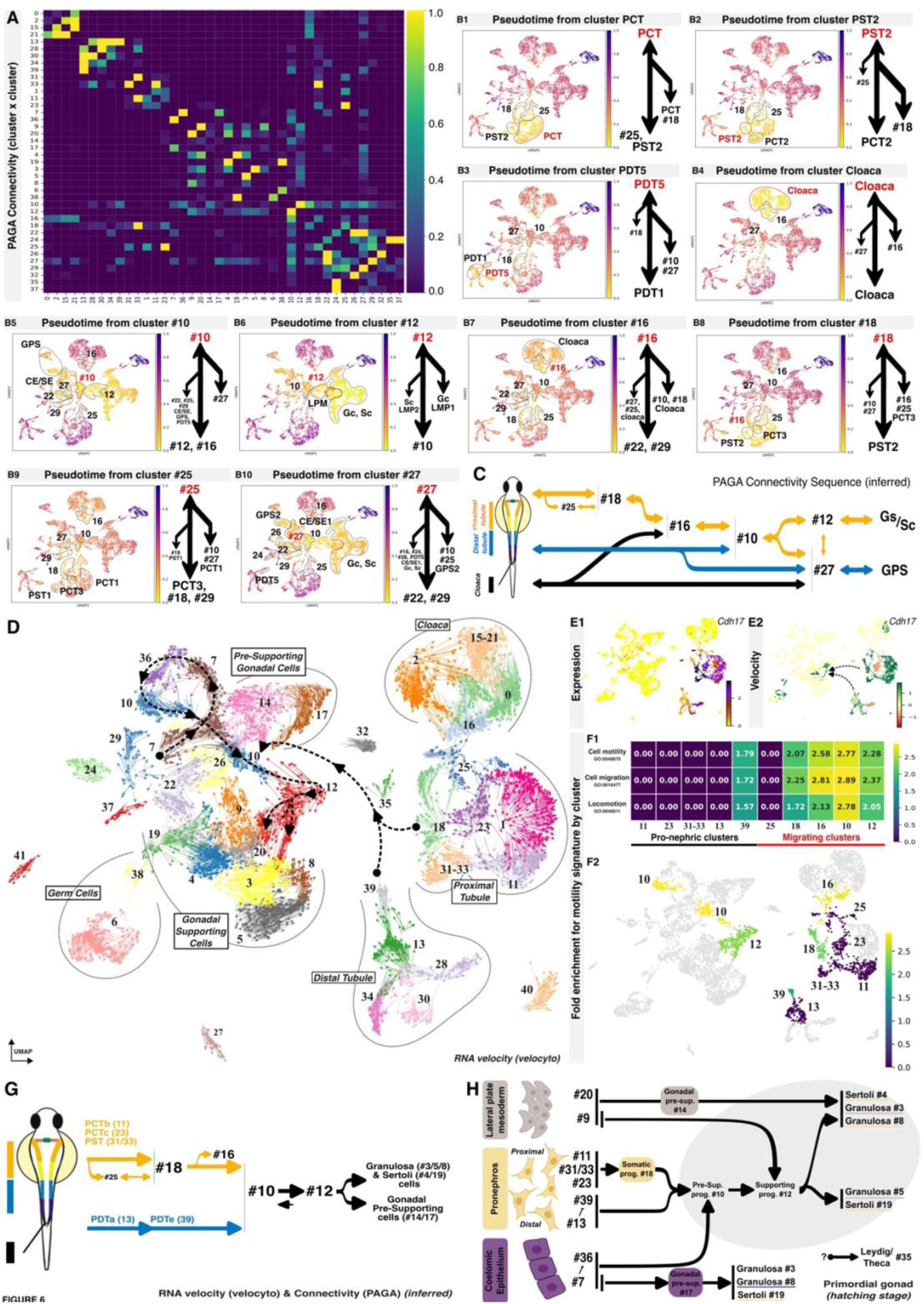
Cell cluster connectivity and trajectory analyses from pronephros to gonad. **(A to H)** Connectivity **(A to C)** and trajectory **(D to G)** analyses were performed across all clusters (as defined in Figure 5). **(A and B)** PAGA-inferred pseudotime indicates connectivity between the different clusters **(A)**. Each value represents the normalized strength of connection based on the fraction of shared nearest-neighbor links between cells of different clusters. Stronger connections indicate a higher degree of transcriptional similarity or transition potential. The color bar indicates connectivity strength, ranging from 0 (no connection) to 1 (maximum connectivity) **(A)**. PCT primarily connects with cluster #25 and Pst2 **(B1)**. PST2 connects mainly with PCT2 and cluster #18 **(B2 and B8)**. PDT5 connects predominantly with PDT1 **(B3)**. Cloaca shows minimal connectivity with other clusters **(B4)**. Cluster #10 connects mainly with clusters #12 and 16 **(B5 and B6)**. Cluster #16 additionally connects with clusters #22 and 29 **(B7)**. Cluster #25 connects primarily with PCT3 and clusters #18 and 29 **(B9)**. Cluster #27 connects mainly with clusters #22 and 29 **(B10)**. Overall, PAGA-inferred pseudotime reveals connectivity between proximal/distal pronephric tubules and cloaca to pre-supporting and supporting gonadal cell clusters *via* cell clusters #25/18/16/10/12/27, #10/12/27 and #27 respectively **(C)**. RNA velocity (velocyto) indicates that cells of pronephric origin (clusters #11, 13, 23, 31/33 and 39) positively contribute to pre-supporting and supporting gonadal fate *via* cluster #10 and 12 **(D)**. Cadherin17 expression **(E1)** and velocity **(E2)** clearly indicate the pronephric contribution to gonad **(E1 and E2)** although expression and velocity are uncorrelated **(E1 compared to E2)**. Motility signatures clearly designate the cells belonging to the pronephros-to-gonad moving clusters (clusters #10/12/16/18/39) having the highest folds enrichment **(F1 and F2)**. Combined PAGA connectivity and RNA velocity analyses allow reconstructing pronephric cell trajectories toward gonadal fate acquisition, primarily *via* clusters #18, #10 and #12 **(G and H)**.

To further elucidate the dynamics of these cell-fate transitions, we next established pairwise PAGA connectivity between the clusters (Figure 6A and 6B; Supplemental Figure 5aA to 5aAN). Stronger connections indicate a higher degree of transcriptional similarity or transition potential (Figure 6A and 6B). As a result, PAGA connectivity clearly links pronephric cell clusters with some of the above-described intermediate cell clusters (Figure 6A and 6B1 to 6B4), and intermediate cell clusters are connected to either other intermediate cell clusters (Figure 6B5 to 6B10) or gonadal cell clusters (Figure 6B6 and 6B10). Overall, the PAGA connectivity analysis delineates both linear and branched relationships between specific pronephric compartments, including the proximal and distal tubules and the cloaca, and the developing gonadal supporting cell lineages (Figure 6C).

Beyond cluster connectivity, we next analyzed expression dynamics in order to estimate RNA velocities at single-cell resolution (Figure 6D). Three distinct cell clusters-one from the pronephros proximal tubule (cluster #18) and two from the pronephros distal tubule (clusters #13 and #39)-project beyond the pronephros toward the gonadal clusters (Figure 6D). Among the intermediate cell clusters, a clear stepwise trajectory emerges, starting from cluster #7, progressing through clusters #36 and #10 and terminating-after the split of cluster #12-with gonadal supporting cell clusters (Figure 6D). For example, while *cdh17* expression is fading outside of the pronephric proximal and distal tubule clusters, trajectory analyses indicate a clear projection toward intermediate cell clusters (clusters #9 and #12; Figure 6E1 and 6E2), demonstrating a progressive fate transition.

To further characterize the pronephric cells migrating towards the gonadal primordium, we next assessed which clusters display motility signatures (Figure 6F1 and 6F2, and Supplemental Figure 6A and 6B). Notably, pronephric clusters #10, 12, 16, 18 and 39 display a motility index well above the average (Figure 6F1 and F2). Among these, only #10 exhibits enrichment for stemness signature (Supplemental Figure 6C and 6D).

Integrating connectivity, developmental trajectories, and motility index not only confirms the developmental links between specific sub-populations of pronephric and gonadal supporting cells, but also maps the stepwise molecular fate underlying their active contributions (Figure 6G).

Different subsets of cells belonging to three cell clusters of the differentiated proximal part of the pronephros (clusters #11, 23 and 31/33; Figure 6D, 6G) project onto cluster #18. Cells within these three clusters retain a robust pronephric contribution characterized by the expression of *cdh17* marker (Figure 5G). Conversely, they lack the expression of gonadal *sox9b* & *GFP* markers (Figure 5H) and fail to exhibit any transcriptional signature associated with cell motility (Figure 6F1 and 6F2). On the other hand, cluster #18 starts to express the sox9b/GFP gonadal marker while residual expression of the pronephric cdh17/mCherry marker is still observed (see Figure 5G and H). Cells of cluster #18 clearly acquire motility (Figure 6F1 and 6F2). Cells of cluster #18 further project onto cluster #10 (Figure 6D). Similarly, cells belonging to the distal part of the pronephros (clusters #13 and 39, Figure 6D) progressively lose their pronephric signature (cdh17/mCherry marker, Figure 5G) and project onto cluster #10. Contributed by clusters #18 and 39, cluster #10 has lost any pronephric signature but acquired a gonadal one (Figure 5G compared to 5H). Cluster #10 exhibits a high motility signature (Figure 6F1 and 6F2) and clearly projects onto cluster #12 as a result of an apparent transitory/transient state. Being very dynamic too, cells of cluster #12 split into two routes, projecting onto gonadal supporting (clusters #3 and 5) and lateral plate mesoderm (cluster #9) clusters (Figure 6D and 6G). Cells belonging to the cloaca (clusters #16 and 21) express the early gonadal markers sox9b and GFP, and are potentially motile (Figures 5H and 6F1 and 6F2), but no specific projections towards gonadal clusters could be determined. The mapping of cell clusters onto gonad-contributing tissues illustrates both divergent and shared differentiation trajectories originating from the LPM, PN, and CE toward pre-gonadal supporting and mature gonadal lineages (Figure 6H).

### Cross-species analysis of pro- and meso-nephric cell contributions to the gonadal molecular landscape

To investigate the evolutionary conservation of the genetic programs underlying transitional states and fate acquisition of the main supporting gonadal cell lineages, we used SAMap [56] (Figure 7A) to identify homologous cell types with shared expression profiles across mice (Figure 7B), chicken (Figure 7C), turtle (Figure 7D) and medaka.

**Figure 7.**
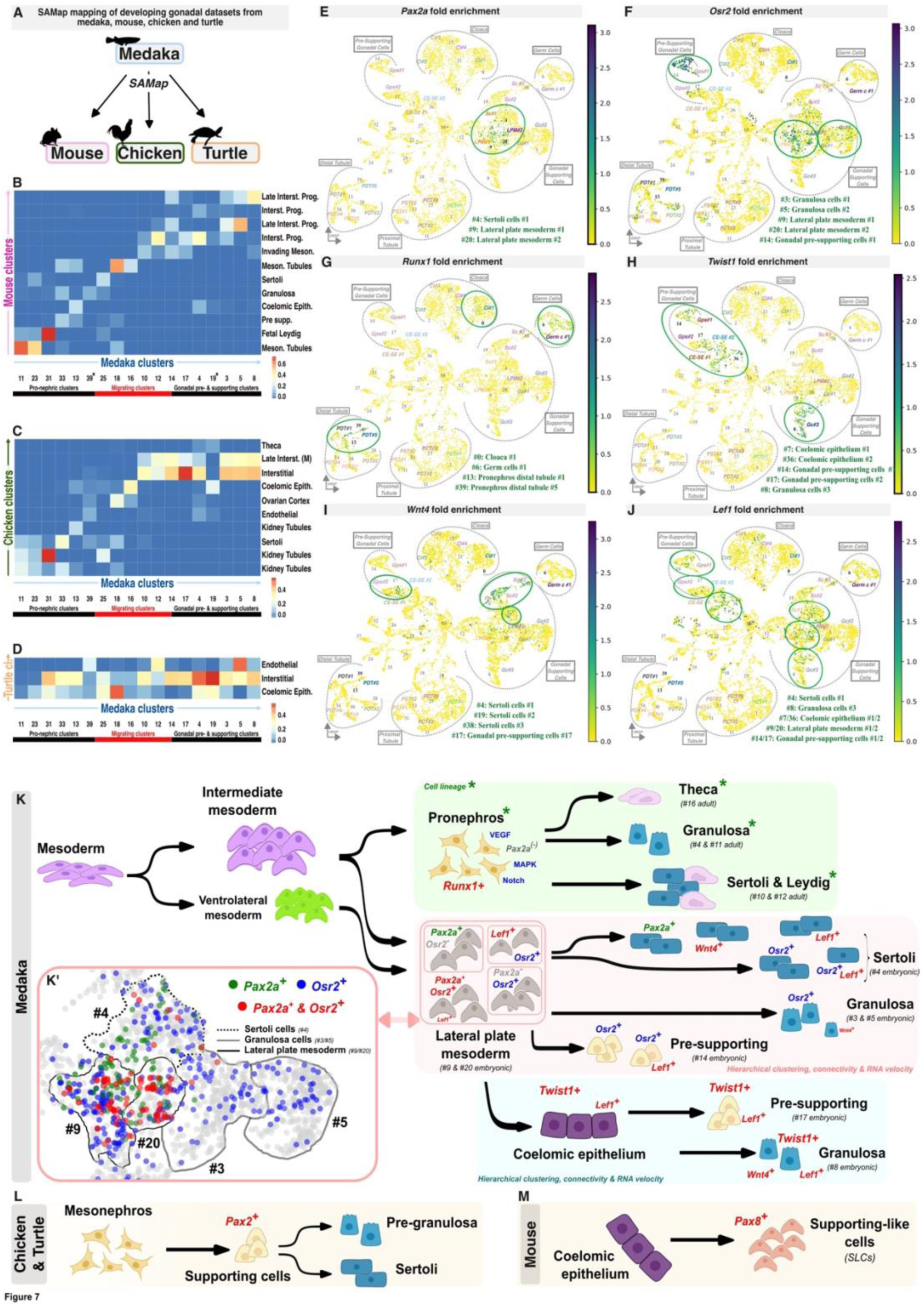
Comparative evolutionary analysis of the molecular landscapes of pro-/meso-nephric cells contributing to gonadal ontogenesis between mice, chicken and medaka. **(A)** SAMap mapping of developing datasets from medaka, mouse, chicken and turtle. **(B to D)** Heatmaps showing Eigengene (ME) scores of medaka pronephric-contributing, migrating and supporting clusters in each of the mouse (**B**), chicken (**C**) and turtle (**D**) gonadal primordium clusters. High ME scores imply that the genes of the medaka clusters are active in the corresponding clusters of either mouse, chicken or turtle. **(E to J)** UMAP reductions of medaka single-cell data for fold enrichment in key genes (*pax2a* (**E**), *osr2* (**F**), *runx1* (**G**), *twist1* (**H**), *wnt4* (**I**) and *lef1* (**J**); relevant clusters are indicated) previously identified as essential for CE/MN cell contribution in mice, chicken or turtle. **(K)** Summary of gonadal soma ontogenesis in medaka, with focus on *pax2a*/*osr2*-expressing cells in the LPM and derived supporting lineages (**K’**). **(L and M)** Cells lineage derivation and progression of *pax*-expressing progenitors in chicken/turtle (**L**) and mouse (**M**) to be compared with medaka (**K**). LPM: Lateral Plate Mesoderm; CE-SE: Coelomic/Supporting Epithelium; Gc: Granulosa cells; Sc: Sertoli cells; Gps: Gonadal pre-supporting cells; Germ c: Germ cells; PDT: Pronephros Distal Tubule; PST: Pronephros Straight Tubule; PCT: Pronephros Convoluted Tubule; Cl: Cloaca.

During the pronephric-to-pre-supporting transition, SAMap alignment scores between medaka and mouse indicate moderate homology for both origin and destination clusters (#11, #23, #31 and #4, #5, #8 respectively) while for transitional states, including migrating clusters, homology clearly decreases (Figure 7B). Comparison between medaka and either chicken or turtle yielded mainly low SAMap alignment scores across all stages, from origin to destination, except for interstitial cell clusters (Figure 7C and 7D) which display high similarity. In both chicken and turtle, single endothelial cell clusters encompass most of the medaka pronephric-to-pre-supporting medaka clusters.

However, higher SAMap alignment scores are observed between medaka, mouse and chicken for the pronephros straight tubule (cluster #31) and granulosa cells (cluster #8). Pairwise alignments further identified shared signatures between medaka and mouse for the convoluted tubules (clusters #11 and #23) and intermediate clusters #18, as well as between medaka and chicken for intermediate cluster #10 (Figure 7B to 7D, and Supplemental Figure 7A to 7C). Comparison of expression levels for key pathways documented to be involved in pronephric-to-pregonadal transitions in mice and chicken (Figure 7E to 7J and Supplemental Figure 8A to 8C) revealed that only the expression of the wnt, notch and vegf pathways are possibly conserved between medaka and mouse. In addition, the hedgehog, TGF-β and ECM pathways emerge as medaka-specific up-regulated pathways during the transitional state (Supplemental Figure 8D and 8E). Similarly, *wnt5*, *fzd*, *plcb*, *patched*, *hhip*, *bcl2*, *gdf6*, *id*, *dcn* and *rspo*, *gli*, *patched*, *pik3*, *bmp4*, *thbs1* are notably up-regulated during either CE or LPM to pregonadal transitions respectively (Supplemental Data 7).

In chicken and turtle, the supporting lineages originate from mesenchymal cells expressing the transcription factor Pax2 [55,57]. In medaka, *pax2a* expression is restricted to the LPM and a sub-population of Sertoli cells (clusters #9, #20 and #4 respectively in Figure 7E). No other *pax* paralogs, including *pax5* and *pax8*, are detected during primordial gonad formation in medaka (Supplemental Figure 8F and 8G). Interestingly, while LPM-*pax2a*-expressing clusters (#9/#20) also transcribe *osr2*, mirroring the mesenchymal expression observed in chicken and turtle [55,57], deeper single-cell analysis revealed additional heterogeneity within these clusters itself. Cells were found to express *pax2a*-only, *osr2*-only or both (Figure 7E and 7F, and Supplemental Figure 9). Upon further differentiation, cells of granulosa clusters #4 & #5 express either *osr2* exclusively or none of these markers, whereas the Sertoli cluster #4 is composed of five distinct sub-populations expressing either *pax2a*, *osr2*, *lef1*, *wnt4* or combinations (Figure 7E to 7J). Regarding expression of other phyla-specific lineage regulators across vertebrates, we could not detect any expression for neither Tbx1 nor Esr1. While *runx1* expression is detected in cloaca, germ cells and pronephros distal tubules (clusters #0, #6, and #13 and #39 respectively in Figure 7G), its direct *ryr2*, *mettl24* or *mmp15* targets are either not expressed or ubiquitously transcribed. Finally, *twist1*, a transcription factor so far only associated with early ovarian differentiation in turtle is highly expressed in CE, gonadal supporting and granulosa cells (clusters #7 and #36, #14 and #17, and #8 respectively in Figure 7H). Altogether, these observations highlight both conserved and derived species-specific variations in the molecular signatures underlying gonadal lineage specification amongst vertebrates.

## Discussion

Our findings challenge the prevailing view that the pronephric field’s contribution to gonadal formation is an amniote-specific innovation. Instead, we show that this trait is not derived but ancestral, predating the divergence of teleosts and tetrapods. We could demonstrate that the teleost gonad is multilineage in origin. However, in contrast to amniotes, the teleost gonad arises from three distinct embryonic tissues: (***i***) the pronephros, (***ii***) the coelomic epithelium, and (***iii***) the lateral plate mesoderm. We found that progenitors derived from these distinct embryonic contributing-tissues give rise to an unexpected lineage-based diversity of adult supporting cell subtypes, including Sertoli and granulosa cells. Equivalent datasets from mouse, chicken and turtle, allowed to decipher whether lineage-specific deviations or adaptive inflections are retained during early gonadal development and the underlying gene regulatory networks.

### Pronephric field contribution is essential for proper gonadal formation and ontogenesis in teleosts

In medaka the apparent permeability between the gonadal primordium and the pronephric anlagen is not merely reflected in the synexpression of marker genes under physiological conditions but also manifests during heat-induced sex-reversion [40]. Wt1 is a key factor in mammalian gonad development where it is expressed early in the urogenital ridge [58,59]. In medaka *wt1a* is first expressed in the LPM during early embryogenesis and later on in the somatic cells of the gonadal primordium. Its paralog *wt1b* is expressed later during embryogenesis but not in the gonadal primordium [60]. We have previously shown that modulating *wt1a* and *wt1b* expression levels impacted the formation of the pronephric field in medaka [60], similar to zebrafish [61,62], linking gonadal primordium development to pronephros formation. Temperature stress applied to medaka embryos does not only result in sex reversion after an illegitimate premature induction of the *dmrt1a* gene in the supporting cells of the primordial gonad but also provokes its ectopic expression in the posterior end of the pronephric duct [40].

Altogether, the analogies amongst expression patterns of supporting, steroidogenic, germ and pronephric cell lineages suggest a physiological interplay between pronephros and early gonad primordia development. Into that direction, cell lineage tracing experiments demonstrate the pronephric contribution to the gonads of both sexes in medaka. We show that all portions of the pronephros (cloaca, distal and proximal tubules) are contributing to the gonad. The absence of any enrichment for stemness signatures suggests that the pronephric-to-gonadal contribution occurs through transdifferentiation, rather than *via* an intermediate de-differentiated state. Although the pronephros contributes cells to both ovarian and testicular lineages without apparent bias, the final functional roles of these contributions differ significantly between the sexes. Hierarchical clustering dendrogram of gonadal supporting cell sub-populations failed to identify any lineage-based correlation among cells of pronephric origin in females. In contrast, a robust lineage-based correlation is apparent in males. Furthermore, subpopulations of Sertoli and Leydig cells derived from the pronephros not only contribute to the general pool of supporting cells (akin to their ovarian counterparts) but also retain a molecular memory of their embryonic origin. Beyond expressing specific Sertoli or Leydig cell markers, these two populations do co-express a physiologically consistent set of genes related to RNA processing. Spatial sub-type-specific repartition of these populations within testis clearly accredits Sertoli and Leydig sub-functionalization according to their origin, especially when of pronephric origin.

### Coelomic epithelium, lateral plate mesoderm and pronephros contribute the gonadal somatic cell lineages in medaka

In mouse, the coelomic epithelium produces supporting, steroidogenic (Leydig/thecal) and non-steroidogenic interstitial cells, while the mesonephric mesenchyme contributes only to steroidogenic lineages (see [2] for review). In contrast, chicken supporting and steroidogenic lineages both originate from mesonephric mesenchyme. Only non-steroidogenic interstitial cells arise from the coelomic epithelium (see [2] for review and [55]).

In medaka the gonadal primordium has been previously described to be possibly derived from two distinct somatic cell populations [22]. While some somatic gonadal precursors arise specifically from the most posterior part of the *ftz-f1* expression domain within the lateral plate mesoderm during early segmentation stage [22], another population, located more dorsally, derives from *sox9b*-expressing cells [48]. Analyses conducted at hatching stage, now provide a single-cell resolution view of primordium gonadal formation in the medaka embryo. Hierarchical clustering established a clear relationship between sub-populations of gonadal/pre-gonadal supporting cells and either the CE or LPM. Together with the pronephric contribution described above, our findings show that the gonadal supporting cell lineage arises from at least three distinct embryonic origins, namely the LPM, CE and PN. Analysis of cell lineage derivation and progression indicated that the supporting lineages of both sexes receive contributions from all these three embryonic layers through a shared axis of progression and differentiation. This axis involves pre-supporting and supporting progenitors. Differentiation from the pronephric distal tubules requires an additional step, transitioning through a somatic progenitor state. Notably, direct differentiations were observed from either LPM or CE to Sertoli and granulosa cell lineages.

Whether these distinct routes of differentiation and transitioning states result in different functional subtypes of gonadal supporting lineages remains to be determined. High-resolution clustering enabled to distinguish two Leydig, three granulosa/theca and four Sertoli cell subtypes in medaka adult gonads. Spatial prediction of single cell clusters corresponding to these different subpopulations further demonstrated distinct and specific testicular localizations (dispersed: Sertoli-a; peripheral: Sertoli-b and Leydig-b or more central: Sertoli-c and –d) according to their subtypes as defined. While these sub-types may represent distinct differentiation states within the same lineage, such patterns could also reflect lineage-specific sub-functionalization of Sertoli and Leydig cells according to their developmental origin. The latter hypothesis appears to hold true for the Sertoli-b and Leydig-b subtypes, which are of pronephric origin whereas the other Sertoli and Leydig subtypes are not. These subtypes, particularly those of the Sertoli/Leydig lineage derived from the pronephros, are likely involved in distinct physiological roles, as evidenced by their involvement in RNA processing.

### The steroidogenic lineage: evolutionary heritage and species-specific adaptations

The developmental source of the fetal and adult steroidogenic cell lineages has long been debated and still remains largely unresolved across mammals and other vertebrates. Recently cell lineage tracing and single-cell transcriptomic analyses supported either dual CE and mesonephric or mesonephric mesenchymal origins in mice and chicken/turtle respectively [55,57,63].

In medaka, single-cell transcriptomics indicated that two subtypes of adult steroidogenic cells, Leydig-b and theca-a, are derived from the pronephros. Hierarchical clustering of granulosa and theca cells supports a lineage-based correlation between both lineages, although pronephric cell lineage tracing, on the other hand, does not indicate any lineage-based correlation between cells of pronephric origin. In contrast, hierarchical clustering of Sertoli and Leydig cell populations does reveal a robust lineage-based correlation, especially when of pronephric origin. These data support a model in which Sertoli and Leydig cell subpopulations derive from a common pool of pronephric progenitors.

During gonad primordium formation, five clusters express genes associated with steroidogenesis: *cyp17* and *cyp17a2* in clusters #4 and #19 which initially display a Sertoli signature, *nr5a1/sf1* in clusters #4, #9 and #19, #20 which exhibit an additional LPM signature, and star in cluster #35. Apparently, while embryonic theca and Leydig cells are not yet clearly specified, embryonic steroidogenesis would nevertheless be mediated by LPM- and Sertoli-committed progenitors as described in turtle and chicken [2,55,57]. However, the presence of an unannotated cell cluster expressing *star*, does not rule out the possibility that interstitial progenitors may also contribute to the steroidogenic cell pool in medaka embryo, analogous to the situation in mouse [54]. In adult gonads, it can be assumed that theca cells are derived from the granulosa pool, while the remaining Leydig cells, which are not of pronephric origin, are derived from the Sertoli cells descending from the lateral plate mesoderm. Nonetheless, although there is no apparent evidence of a steroidogenic cell sub-population derived from the CE, a potential source from the interstitial lineage cannot be excluded. Altogether, our findings point to evolutionary variations in the developmental origins of the steroidogenic cell types among vertebrates.

The derivation of teleost steroidogenic cells from the supporting cell lineage might reflect a developmental program more similar to that of chicken and turtle. However, in teleosts, this population receives an additional contribution from both the LPM and the PN, whereas in chicken the lineage originates exclusively from the mesonephros. Though, the potential contribution –while likely marginal in medaka-of an additional interstitial lineage, comprising *bona fide* fetal/embryonic cells “dedicated” to steroidogenesis, evokes a mammalian-type developmental ontogeny. By contrast, the absence of such cells in chicken [55] and turtle [57] raises questions about the evolutionary trajectories of steroidogenic function and its cellular mediators across vertebrates. These results highlight that the steroidogenic function, while intrinsically conserved on the biochemical level across clades, can be carried out by diverse gonadal cell lineages ultimately converging on similar functional outcomes as adaptive inflections of an otherwise primarily conserved underlying gene regulatory network.

### Gene regulatory networks and evolutionary perspective

In chicken and turtle, the supporting cell lineage originates primarily from Pax2+ mesenchymal cells of the mesonephros [55,57]. In mouse, equivalent supporting cells emerge predominantly from the CE, and not from Pax2+ mesenchymal cells. Nevertheless, a recently identified population of gonadal progenitors arising from the mesonephric-gonadal interface, the supporting-like cells, contributing to the formation of the rete testis and rete ovarii in mammalian gonads and constituting an additional source of Sertoli and granulosa cells, does express *pax8* [64–66], a close paralog of Pax2. In medaka, while *pax8* expression is not detected, both LPM populations that give rise to part of the supporting lineage express *pax2*. It seems reasonable to assume that pax2-expressing cells contributing to the supporting cell lineage represent the ancestral state. However, the shared ontogenetic origin of the pro-/mesonephros and the LPM from a common precursor population within the intermediate mesoderm [67]-marked by the expression of *Lhx1*, *Osr1*, and *Pax2* [68]-suggests a conserved genetic framework underlying the diversification of supporting cell lineages across vertebrates. Though the functional substitution of *Pax2* by *Pax8* during mammalian evolution suggests a shift in transcriptional regulation, the ontogenetic diversity of these gonadal precursor cells (LPM, pro-/meso-nephros or CE, see Figure 7K and 7K’) remains puzzling from an evolutionary perspective. This diversity highlights significant questions regarding the selective pressures and developmental mechanisms driving such lineage divergence.

Answer may come from the observation that specifications of supporting cell lineages across vertebrates is obviously driven more by differences in developmental timing than by fundamental changes in gene expressions or cell-type identity. This implies that heterochronic shifts (alterations in the timing of developmental events) may play a pivotal role in shaping lineage-specific commitment or adaptations, rather than the evolution of entirely novel cell types. Thus, the predominance of the CE as the major source of supporting gonadal cells in mammals, as opposed to the mesonephros in chickens and turtles or the PN/LPM/CE in medaka, could reflect a heterochronic shift in the deployment of the intermediate mesoderm and its nephric derivatives.

The lineage-specific usage of *Wnt4*, *Lef1*, *Lhx1*, *Osr1*, and *Pax2/8* further supports this hypothesis (see Figure 7E to 7M). These transcription factors, conserved across vertebrates, may act as molecular switches that respond to temporal cues during development, directing precursor cells from the intermediate mesoderm toward distinct fates -CE, LPM, or pronephros/mesonephros-depending on the lineage. This perspective underscores the importance of developmental timing as a key driver of evolutionary innovation, offering a unifying framework to explain the diversity of supporting cell lineages across vertebrates.

### In light of evolution: lineage-specific deviations, convergent evolution or adaptive inflections?

Although the overall architectures of testes or ovary are conserved across vertebrates, the underlying ontogenetic processes are far more plastic than previously assumed. While cell migration and ingression appear to be ancestral and conserved features of gonadal primordium assembly and formation, significant divergences exist in the identity of mobilized cells, their respective contributions, the molecular pathways involved, and the transitional states guiding their fate commitment.

The prevalence of the CE as the primary source of supporting gonadal cells in mammals [65,69,70], contrasts with the mesonephros origin in chicken [55] and turtle [57], and even more with the combined PN/LPM/CE contribution in medaka. Into that direction, the greatest similarity of medaka pro-nephric, migrating and gonadal pre-supporting cells clusters to only a few endothelial or interstitial cell clusters in chicken, turtle and (to a lesser extent) in mouse, suggests evolutionary lineage reductions –whether in cell types themselves or in the sequential timing of their differentiation-. If the teleost situation reflects the ancestral system, this implies that the process of gonad development has undergone lineage-specific simplifications during the course of evolution. A similar configuration is observed in early steroidogenic functions, which are mediated by supporting progenitors and fetal split populations in medaka that are either already Sertoli-fate committed or more dedicated (potentially interstitial), whereas in mammals and turtles/birds these functions are endorsed by dedicated fetal or supporting progenitors only. In both contexts, the core regulatory components, such as *pax2a*, *twist1*, *wnt4*, *lef1* or *runx1*, see Figure 7, or steroidogenic factors like *cyp17*, *cyp17a1*, *star* or *nr5a1/sf1*, are similarly co-opted by distinct different progenitor populations during the course of evolution. This is especially evident for the *pax* genes which engage supporting cell lineage commitment from diverse progenitor sources: the LPM in medaka, the mesonephros in turtle/chicken, and the CE in mouse (see Figure 7K to 7M). Interestingly the findings in medaka also reveal that the spatial repartition of adult Sertoli sub-type-specific cell populations clearly accredit an evolutionary-based sub-functionalization according to their embryonic tissue of origin, particularly when of pronephric origin, that has not been described so far in other organisms.

Whether these variations reflect lineage-specific deviations from an ancestral state, convergent evolution or adaptive processes of an overall universal developmental potential driven by physiological constraints toward a similar gonadal organization are definitively the next challenge to be addressed. It is now evident that, over the course of evolution, the different species achieve similar phenotypic outcomes primarily through conserved pathways by co-option and rewiring of pre-existing gene regulatory networks and shifts in developmental timing rather than by identical progenitor cell types. Regardless of the initial cell type executing these networks, functional outcomes remain consistent, as a result of a canalized developmental system drift.

## Supplemental Figures

**Supplemental Figure 1.**
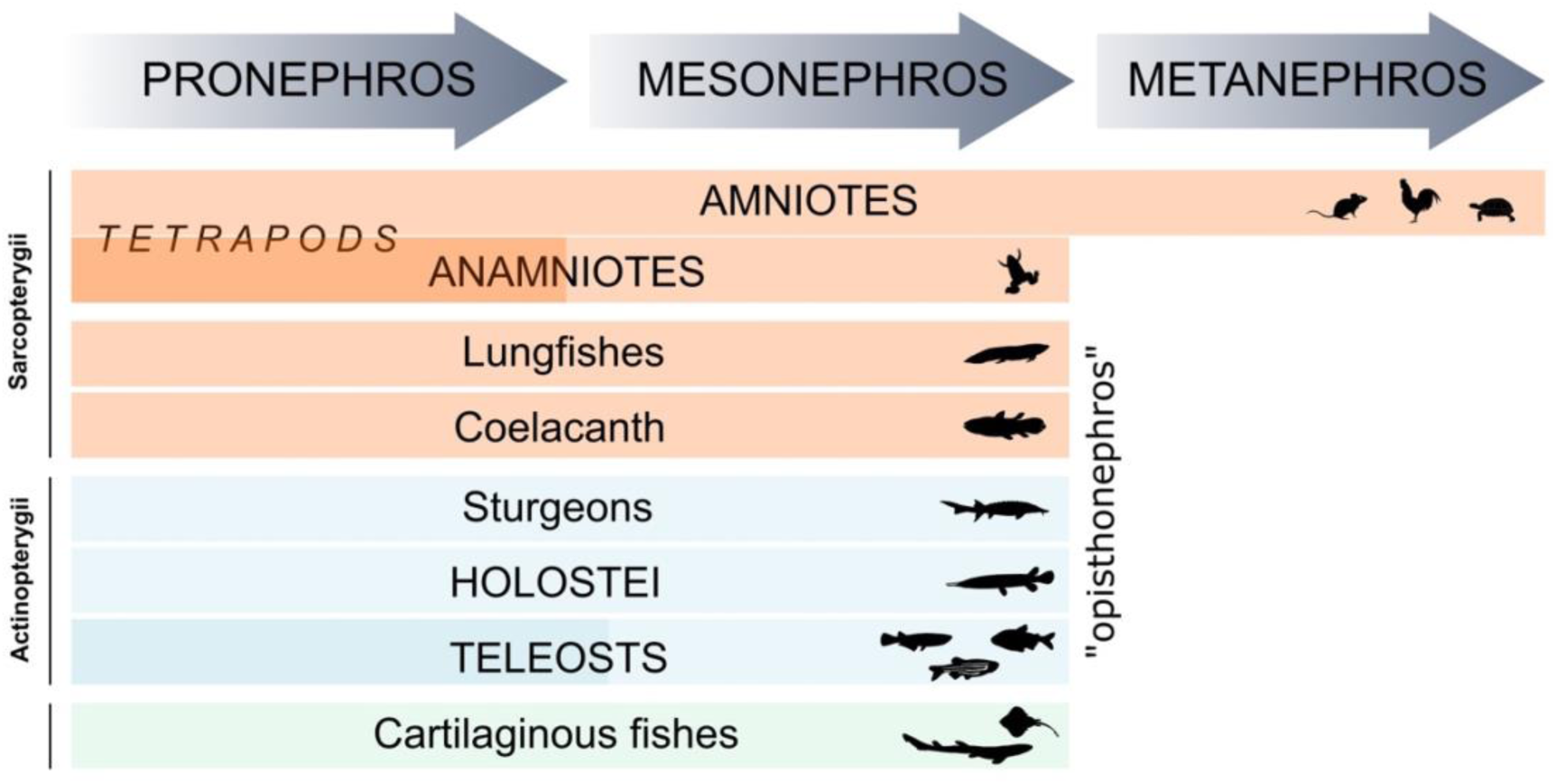
Embryonic kidney development across vertebrates.

**Supplemental Figure 2.**
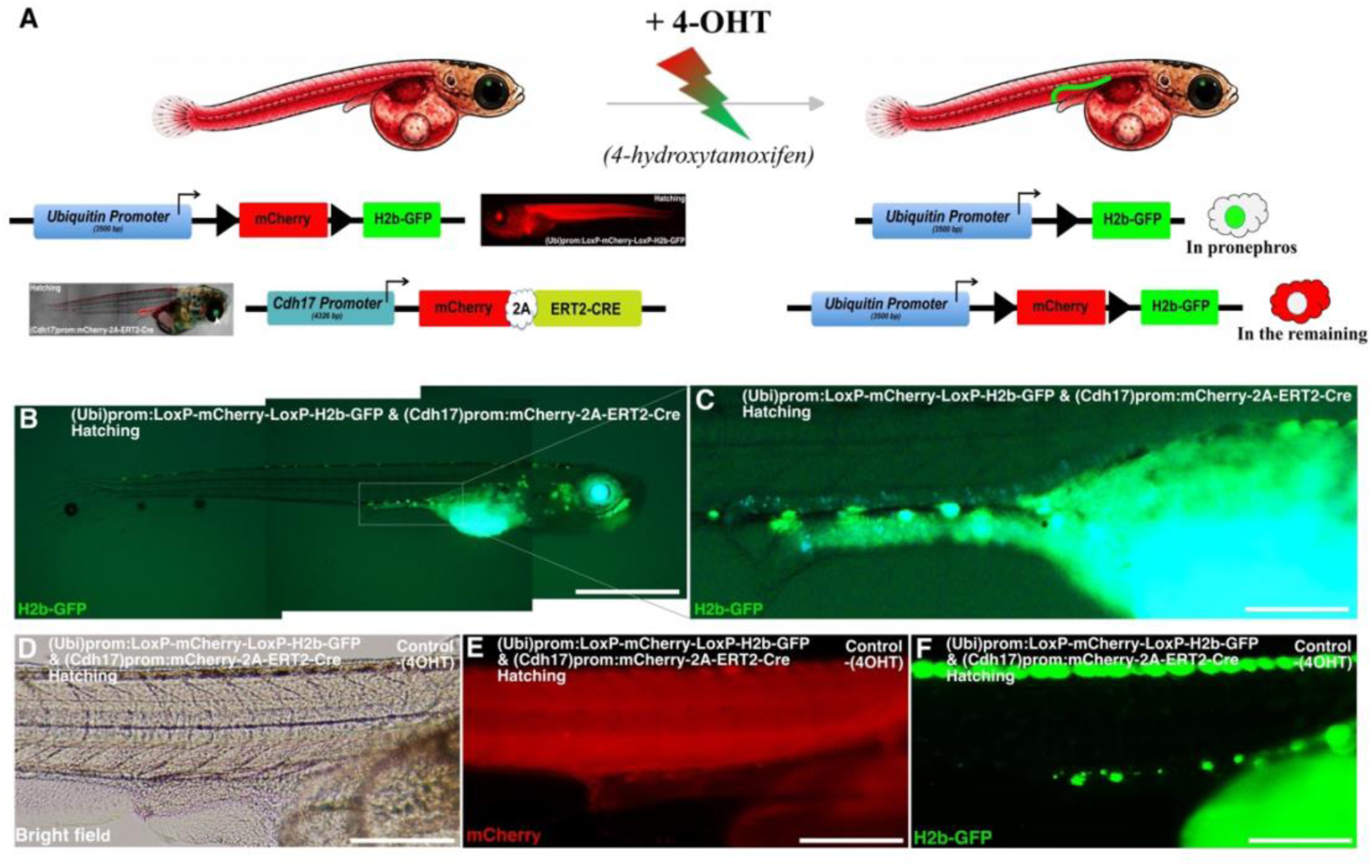
Generation of a *cdh17* medaka model for inducible *in vivo* cell lineage tracing of early pronephric field cells. **(A to C)** Upon 4-OHT exposure, CRE recombinase - expressed under the pronephric-specific *cdh17* promoter - translocates to the nucleus, inducing recombination and excision of the mCherry-stop cassette. This results in permanent GFP labelling of pronephric cells. **(D to F)** In absence of 4-OHT, CRE recombinase remains cytoplasmic, and no recombination (GFP fluorescence) is detected. Scale bars, **B and C**, 450 µM; **D to F**, 200 µM.

**Supplemental Figure 3.**
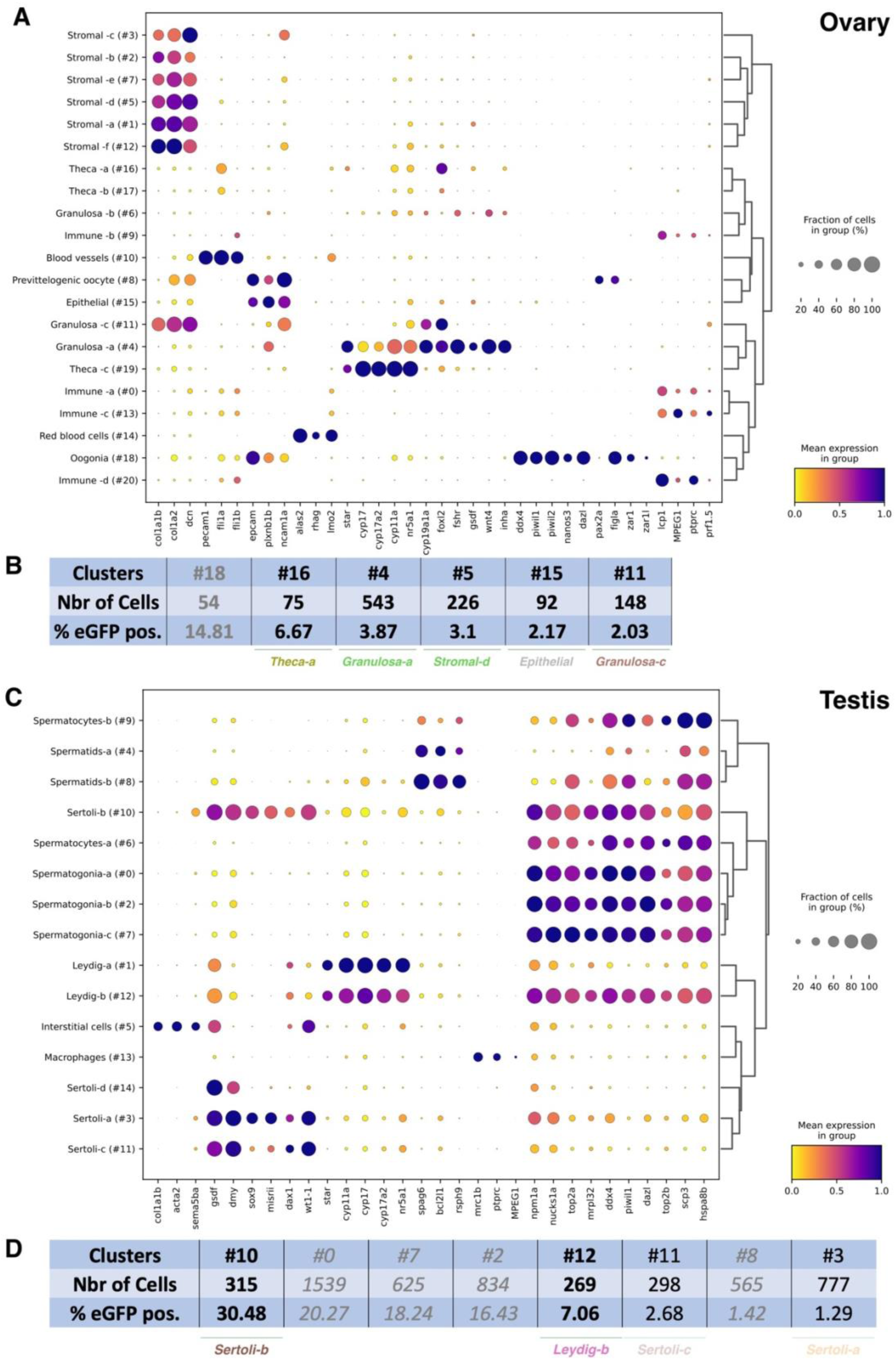
Marker genes and marker transcription factors of the cell types of Medaka adult gonads (testis and ovary). **(A and C)** Dotplots displaying the expression of the main non-redundant markers per leiden cluster. Sizes of the dots represent the percentage of the cells expressing the gene in each cell type and the color the mean expression. (**C and D)** Tables showing the percentages of cells of pronephric origin (GFP-expressing) within each cluster.

**Supplemental Figure 4.**
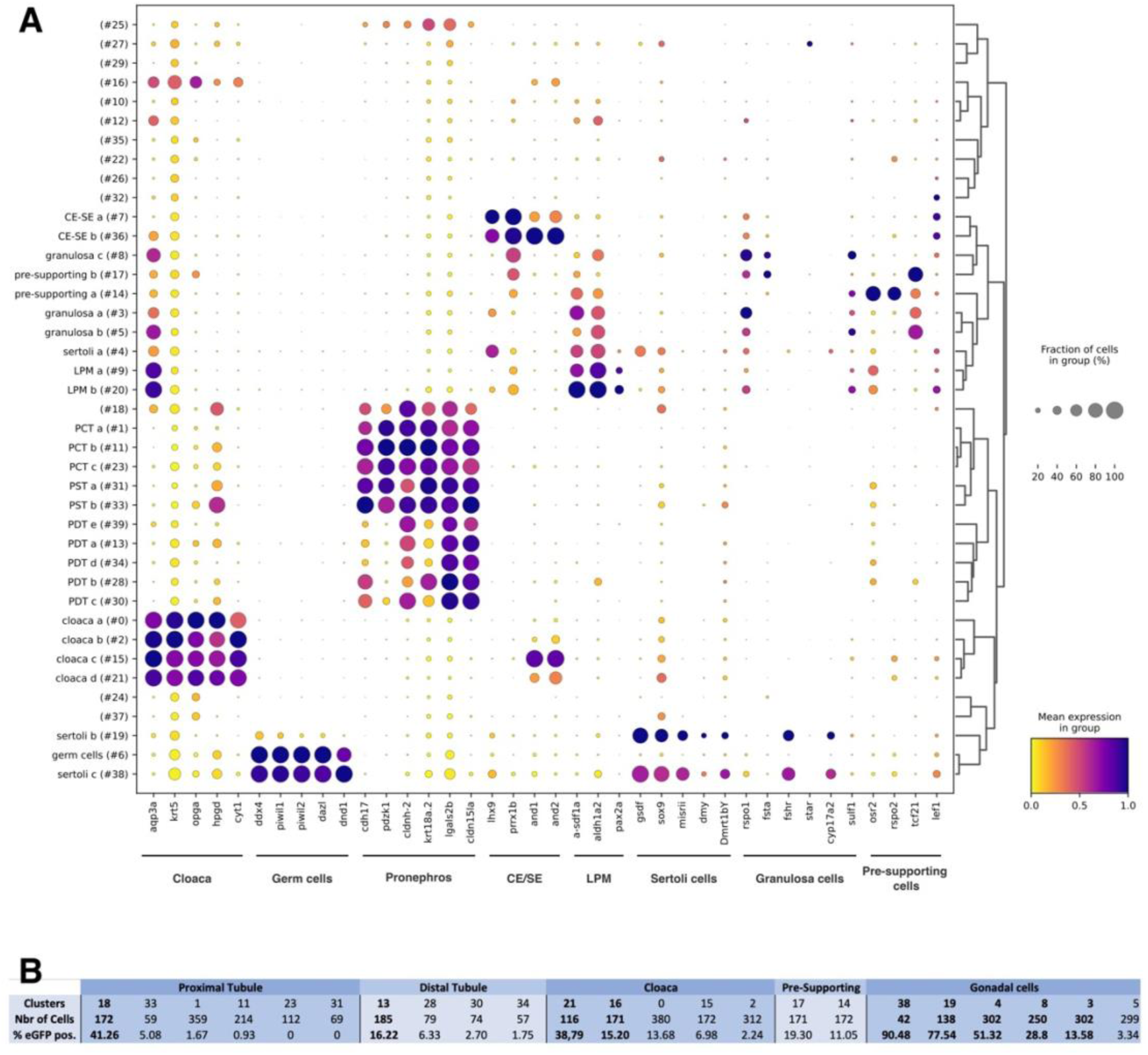
Marker genes and marker transcription factors of the cell types of Medaka primordial gonads. **(A)** Dotplots displaying the expression of the main non-redundant markers per Leiden cluster. Sizes of the dots represent the percentage of the cells expressing the gene in each cell type and the color the mean expression. (**B)** Percentages of cells of pronephric origin (GFP-expressing) within each cluster.

**Supplemental Figure 5.**
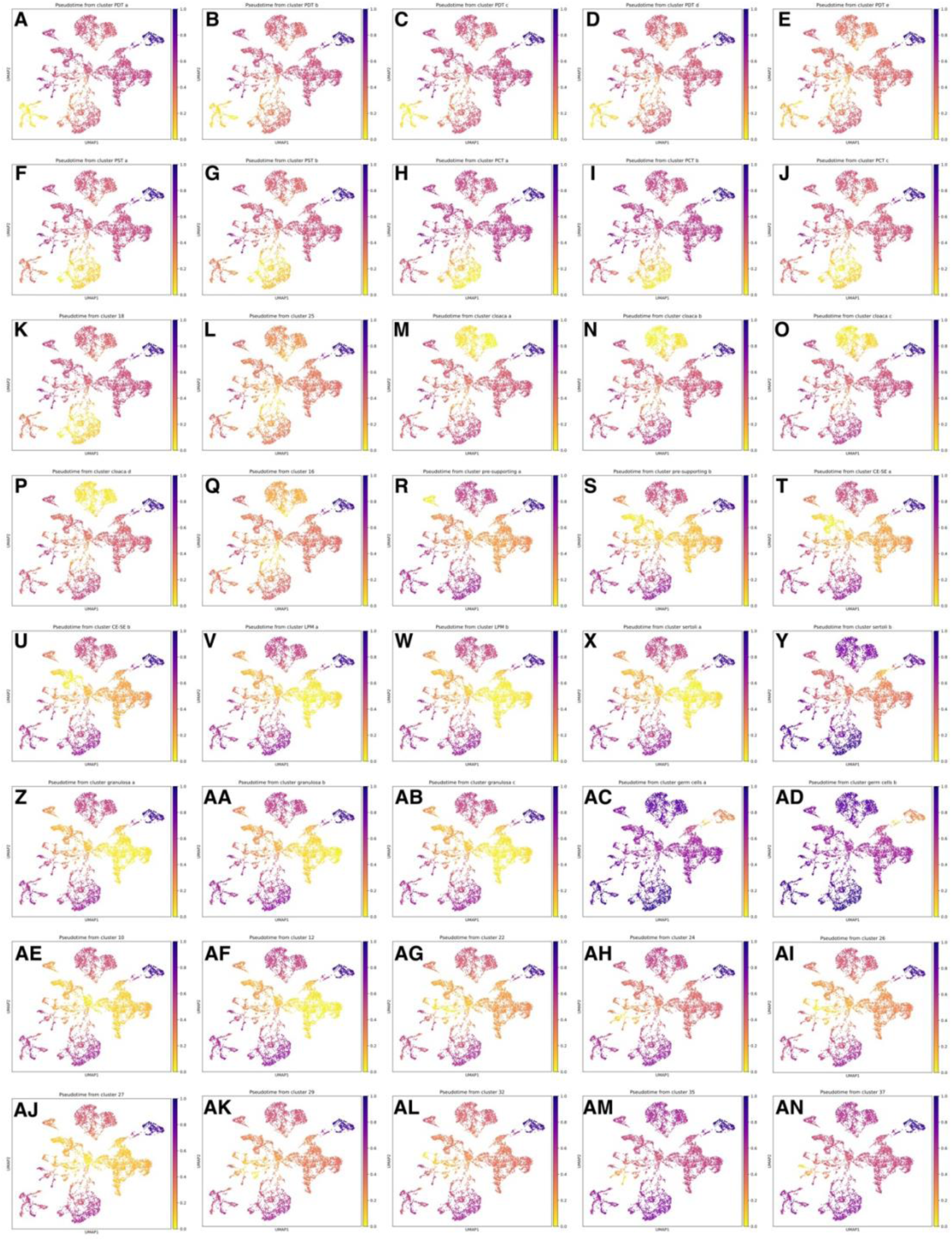
Cell cluster connectivity analyses from pronephros to gonad. **(A to AN)** Connectivity analyses were performed across all clusters (as defined in Figure 5). PAGA-inferred pseudotime indicates connectivity between the different clusters. Each value represents the normalized strength of connection based on the fraction of shared nearest-neighbor links between cells of different clusters. Stronger connections indicate a higher degree of transcriptional similarity or transition potential. The color bar indicates connectivity strength, ranging from 0 (no connection) to 1 (maximum connectivity).

**Supplemental Figure 6.**
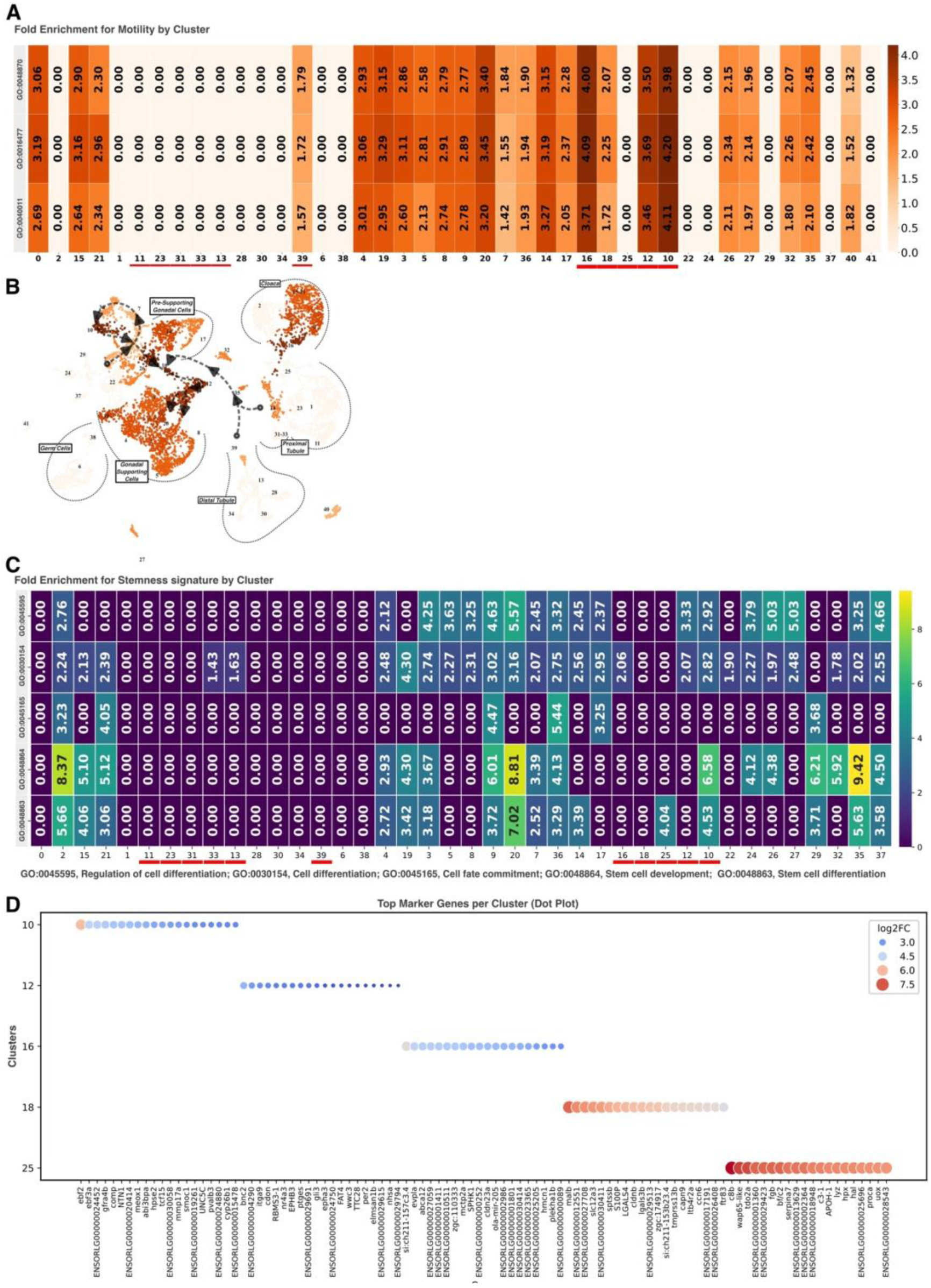
Fold enrichment for motility (A and B) and stemness (C) signatures and top marker genes by clusters encompassing migrating cells (D) during pronephric-to-pre-supporting gonadal transition in medaka embryonic gonads.

**Supplemental Figure 7.**
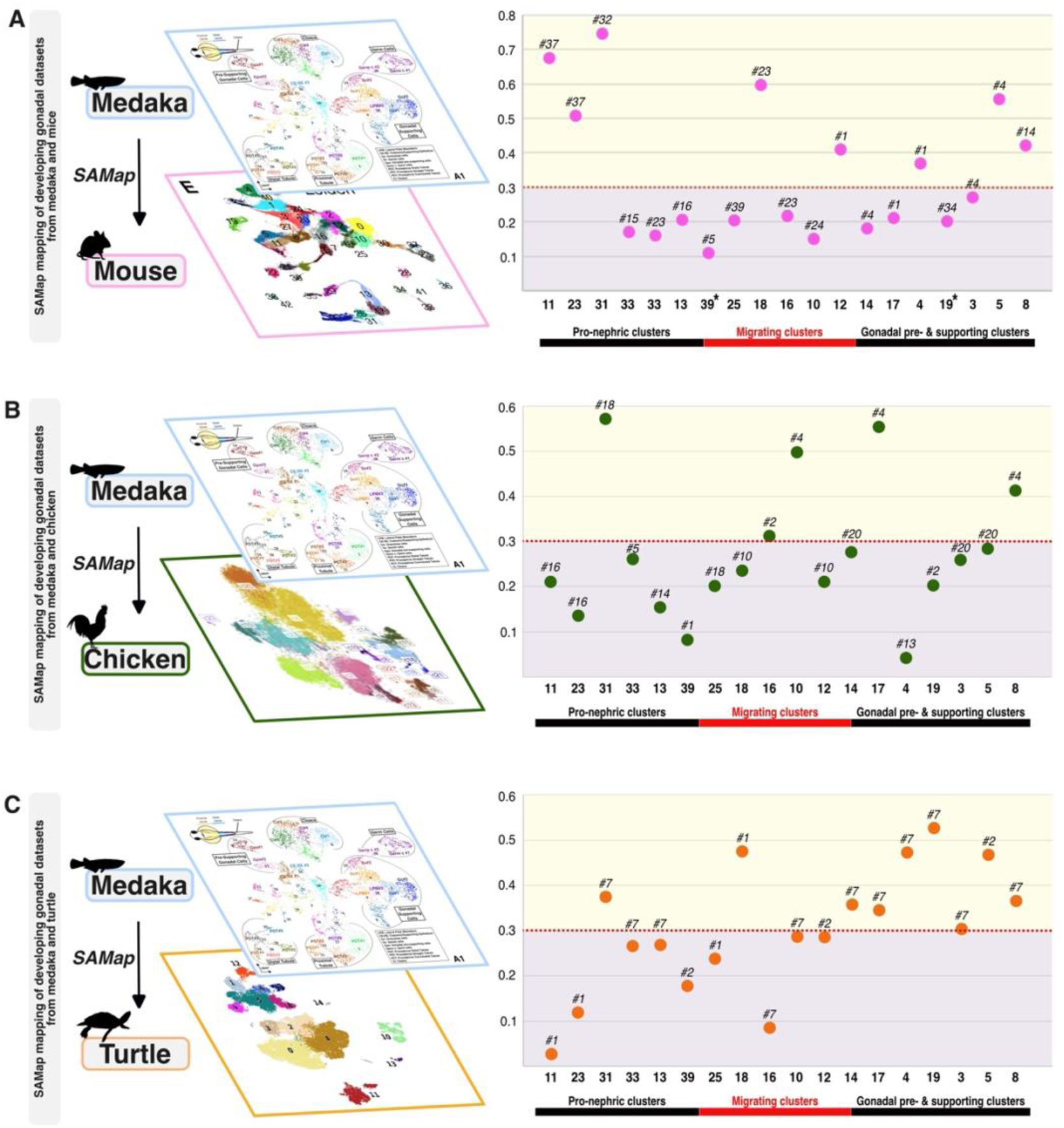
Comparative evolutionary analysis of the molecular landscapes of pro-/meso-nephric cells contributing to gonadal ontogenesis between mice, chicken and medaka. **(A to C)** Homologous cell types (clusters, Eigengene (ME) scores) with shared expression programs between medaka and either mouse **(A)**, chicken **(B)** or turtle **(C)**. High ME scores imply that the genes of the medaka clusters are active in the corresponding clusters of either mouse, chicken or turtle.

**Supplemental Figure 8.**
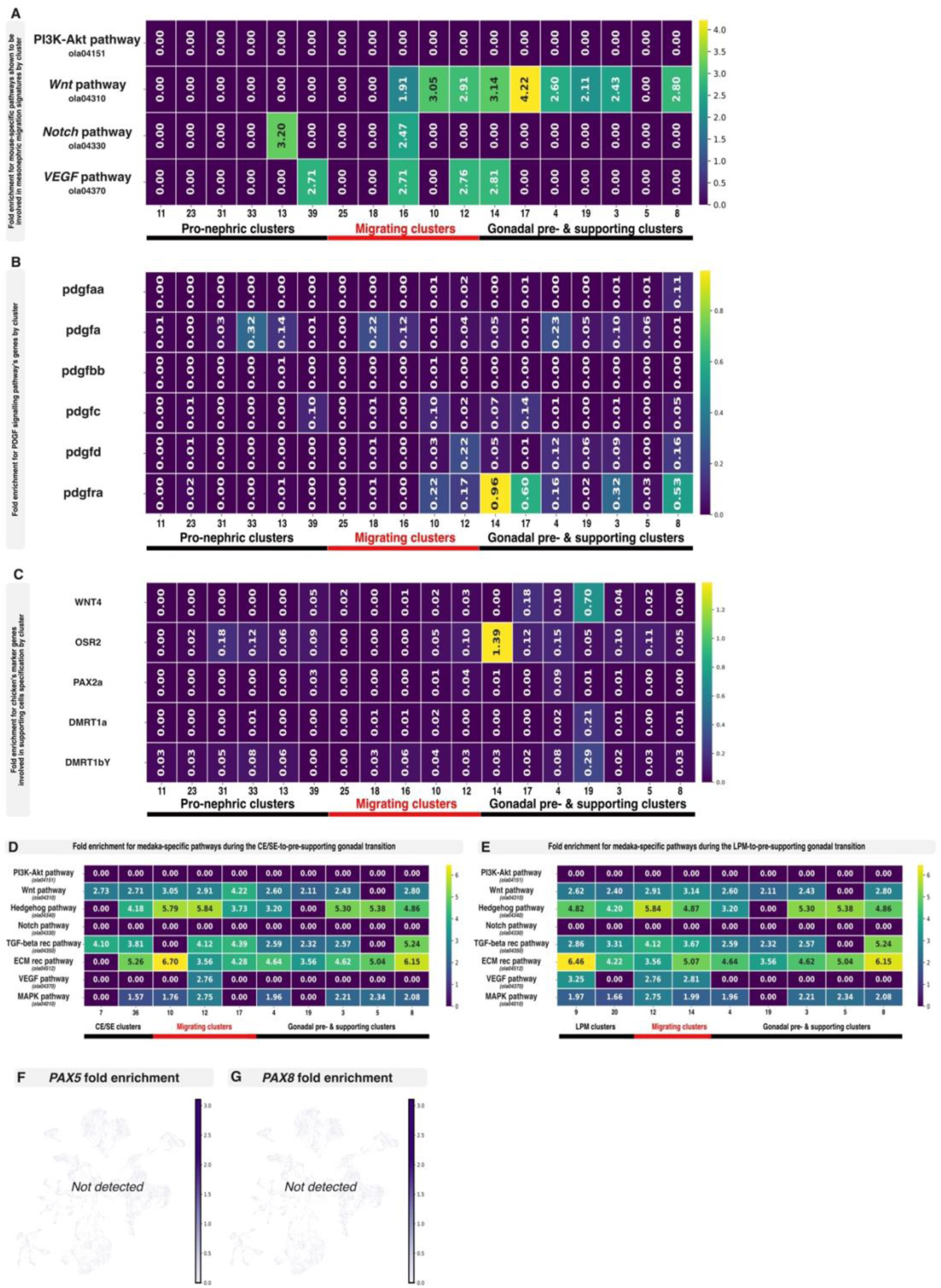
Signalling pathways fold enrichment. Heat map of fold enrichment in key pathways (PI3K-Akt, Wnt, Notch and VEGF) previously identified as essential for CE/MN cell contribution in mice **(A and B)** and chicken (Wnt4, Osr2, Pax2a, Dmrt1a, Dmrt1bY, **(C)**). Heat map of fold enrichment in key pathways (PI3K-Akt, Wnt, Hedgehog, Notch, TGF-beta, ECM, VEGF and MAPK) during either CE/SE-to-pre-supporting or LPM-to-pre-supporting transitions in medaka primordial gonads **(D and E)**. UMAP reduction of Pax5 **(F)** and Pax8 **(G)** expressions during primordial gonad formation in medaka.

**Supplemental Figure 9.**
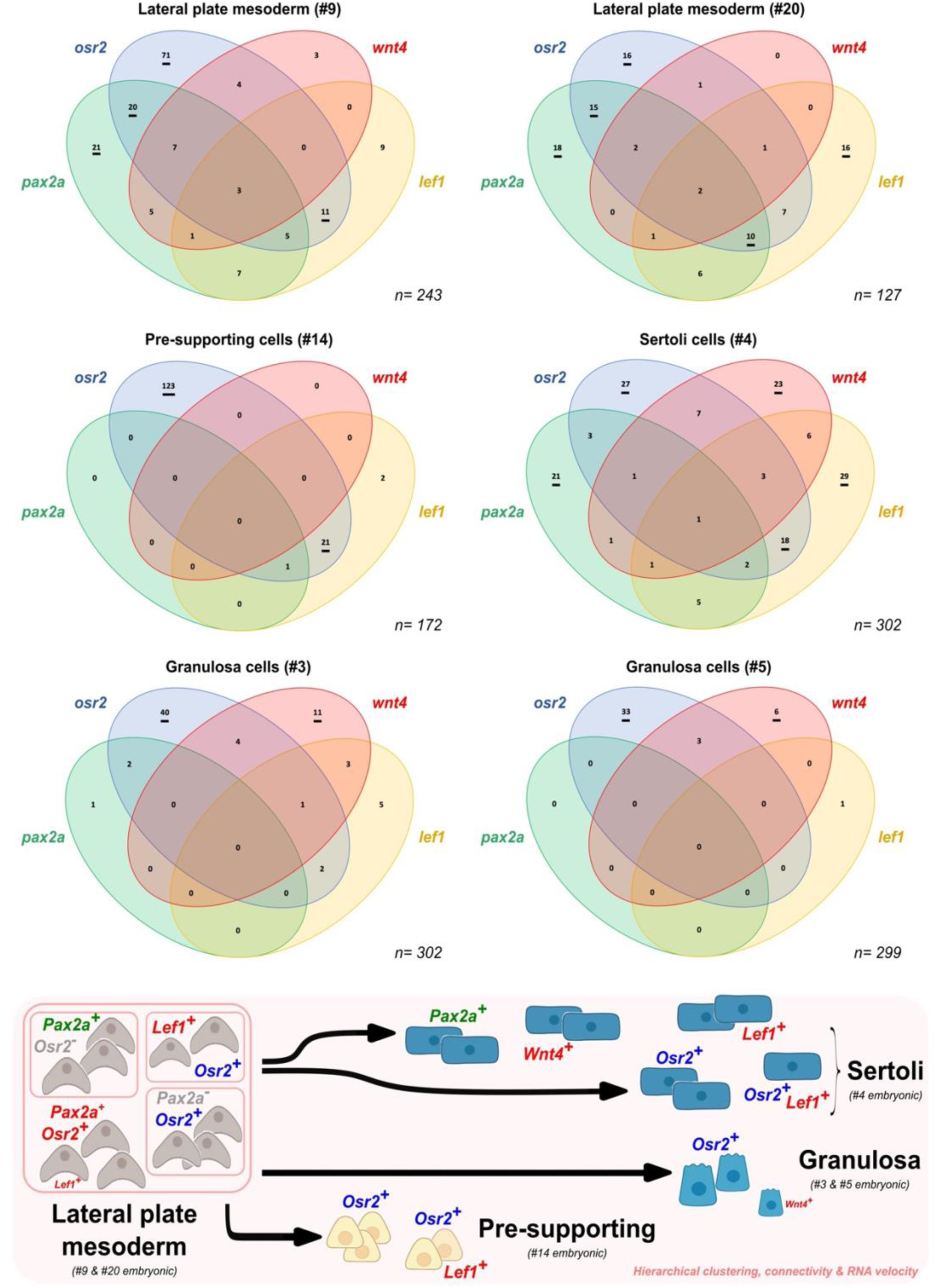
Cells lineage derivation and progression of *pax2a*-expressing progenitors in medaka. Analysis of medaka *pax2a*-derived cells from the lateral plate mesoderm reveal an unexpected diversity of sub-populations of cells expressing different combinations of marker genes, *pax2a*, *osr2*, *lef1* and *wnt4* for instance. See also Figure 7.

## STAR Methods

### Resource availability

#### Lead contact

Further information and requests for resources and reagents should be directed to and will be fulfilled by the lead contact, Amaury Herpin.

#### Material availability

Unique reagents generated in this study are available from the lead contact with a completed Material Transfer Agreement.

#### Data and code availability

Raw single-cell RNA-seq data have been deposited in the European Nucleotide Archive (ENA) under accession number PRJEB108462. Other relevant data and materials, as well as any additional information required to reanalyze the data reported in this paper, are available from the lead contact upon request.

### Experimental model

Medaka fish used in this study were taken from closed breeding stocks of the HdrR *Oryzias latipes* strain (WLC 3860). The animals were kept under under standard photoperiod cycle of 14 hr/10 hr light/dark at 26°C (+/-1°C). Eggs were collected 1-2 hr after starting the light period and raised at 25°C in Danieau’s medium (17.4 mM NaCl, 0.21 mM KCl, 0.12 mM MgSO_4_, 0.18 mM Ca(NO_3_)_2_, 1.5 mM Hepes, buffered at pH 7.2). Stages of development were described according to Iwamatsu [37]. Animals were kept and sampled in accordance with the applicable EU and national French legislation (EU Directive 2010/63/EU) governing animal experimentation and protection.

### Method details

#### Microinjection

Medaka one-cell stage embryos were injected as described [39,74]. In details, using glass capillary needles and a FemtoJet device (Eppendorf), a DNA solution (5 to 30 ng/µL) is injected (about 1 nL volume per cell) into the first blastomere shortly after fertilization (5 to 20 min after fertilization). For injection embryos were kept in 1x Danieau’s solution at 18°C, fixed in an agarose mold. Injected embryos are then transferred to a 27°C incubator chamber for subsequent development.

#### Establishment of transgenic fluorescent reporter lines and imaging

For a dynamic and *in vivo* visualization of endogenous marker genes expression several transgenic fluorescent reporter lines were employed. ***Ol-bsf/LRPPRC*** see [75]; ***9Kb-dmrt1bY*-promoter** see [76]; ***Ol-follistatin*** see [77]; ***sox9b*** see [48], ***Ol-vasa*** see [78] and ***dmrt1a-GFP*** see [40]. For ***cadherin17-mCherry***, the upstream promoter region of the *Ol-cadherin-17* gene (4.325 bp upstream the transcription start) was cloned (*Xba1* sites) in front of the mCherry ORF of a meganuclease plasmid. For the generation of a stable transgenic line, the meganuclease protocol was used. Briefly, approximately 10 to 15 pg of total vector DNA in a volume of 500 pL injection solution containing *I-Sce1* meganuclease was injected into the cytoplasm of one-cell-staged medaka embryos. Adult F0 fish were mated to each other, and the offspring were tested for the presence of the transgene by fluorescence check. Siblings from positive F1 generation fish were raised to adulthood and tested again for fluorescence. For imaging, embryos, hatchlings or tissues were slightly fixed (4% paraformaldehyde for 15 minutes on ice) and then mounted with 1-2% low melting temperature agarose. Confocal pictures and image stacks were acquired using a Nikon C1 (eclipse Ti) confocal laser scanning microscope and the NIS elements AR software.

#### Morpholino microinjection

For Ol-cdh17 knockdown experiments, embryos were injected with an ATG morpholino targeting the transcription start of the cdh17 mRNA (XM_004078115). The most efficient dose (4 mg/mL) was experimentally determined. For progressive impaired development of the pronephros, doses ranging from 0.5 mg/mL to 4 mg/mL were employed.

#### Cell lineage assay (transgenic lines

Two transgenic lines (driver and effector line) were created for the *in vivo* cell lineage assay. (***i***) <u>Driver line</u>: a 4.326 bp cadherin17 promoter region upstream the transcription start was cloned (*Kpn1*/*Cla1*) in front of an [mCherry-2A-ERT2Cre]. (***ii***) <u>Effector line (Gaudi^RSG^, [79])</u>: a 3.5 Kb zebrafish ubiquitin promoter replaced a *Hsp70* promoter in Addgene plasmid 24334 [80], resulting in the pBS/I-Sce1/Ubiquitin::Loxp-DsRed-LoxP-H2B-eGFP plasmid.

For activation of the Cre recombinase and subsequent effective recombination, embryos (hatching stage) were treated with 0.5 µM 4-hydroxytamoxifen (4-OHT; sigma H6278) for three consecutive days. Embryos were kept in the dark throughout the duration of the treatment. At the end, embryos were washed with 1X Yamamoto’s solution.

#### Fluorescence-activated cell sorting (FACS)

Sox9b-GFP and cdh17-mCherry double transgenic hatchlings of medaka were used for this experiment. Approximately 350 gonadal fragments, including both gonads and pronephros, were dissected from embryos at different developmental stages: hatching, 2-3 days post-hatching (dph), 4 dph and 6-7 dph. Cells were dissociated by incubating the gonadal tissue with 0.05% trypsin-EDTA (Fisher Scientific, 11580626) for 15 minutes at room temperature. After dissociation, the cell suspension was transferred to Medium 199 (M199) (Gibco, 22350029) supplemented with 15% fetal bovine serum (FBS) (GE Healthcare, A11-151) to neutralize the trypsin activity. The suspension was filtered through a 40 µm cell strainer to remove clumps and then centrifuged at 200g for 5 minutes. The resulting cell pellet was resuspended in phosphate-buffered saline (PBS) (Sigma, P8537). Sox9b-GFP, cdh17-mCherry, and sox9-GFP/cdh17-mCherry-positive cells were isolated by FACS (FACSAria FUSION, Becton Dickinson) based on fluorescence intensity and cell size. For single cell analysis, cells were labelled with DAPI (1µg/mL). Viable (DAPI negative), Sox9b-GFP, cdh17-mCherry, and sox9-GFP/cdh17-mCherry-positive cells were then isolated using a standard MACSQuant Tyto Cartridge on MACSQuant® Tyto® (Miltenyi Biotec). Cell viability, sorting purity, and numeration of the sorted fraction were evaluated on a MACSQuant Analyzer 16 (Miltenyi Biotec) before single cell library preparation.

#### Single-cell capture, library preparation, and sequencing

Single-cell suspensions were obtained from 2-3 dpf embryos, testis, and ovary using optimized enzymatic dissociation protocols to preserve transcriptomic integrity. Cell viability (>90%) was confirmed prior to encapsulation. Cells were loaded onto the Chromium Single Cell 3′ platform (v3 chemistry, 10x Genomics), where individual cells were partitioned into Gel Bead-In-Emulsions (GEMs) for barcoding and reverse transcription. Libraries were prepared according to the manufacturer’s protocol and sequenced on an Illumina NovaSeq X Plus platform. Sequencing depth varied across libraries, with a median of 36,000 reads per cell (range: 28,000-41,000). **Pre-processing and quality control** Raw sequencing data were processed using Cell Ranger (v7.1.0), including demultiplexing, alignment to the *Oryzias latipes* genome assembly ASM223467v1 (Ensembl release 107), and UMI counting. Downstream analyses were performed using Scanpy (v1.10.4). Genes detected in fewer than 10 cells were excluded. Cells with fewer than 200 detected genes or fewer than 500 total counts were removed. Additional filtering excluded cells with mitochondrial transcript fractions greater than 5% or ribosomal transcript fractions greater than 30%. Putative doublets were identified and removed using Scrublet. Only high-quality singlet cells were retained for downstream analyses. Counts were normalized per cell to equalize total library size and log-transformed. Highly variable genes were identified using the Seurat v3 flavor of variance-stabilized dispersion selection. Following QC, filtering, and doublet removal, altogether 6.321, 9.656, and 8.094 cells were sequenced for hatchlings, male and female adult gonads respectively.

#### Dimensionality reduction, neighbourhood graph construction, clustering, and annotation

Data were scaled and subjected to principal component analysis. The resulting principal components were used to construct a k-nearest neighbor graph based on Euclidean distances in PCA space (n_neighbors = 14). Cell clustering was performed using the Leiden algorithm with a resolution parameter of 0.3. Two-dimensional embeddings were generated using Uniform Manifold Approximation and Projection (UMAP) with parameters min_dist = 0.4 and spread = 1.0, computed on the k-nearest neighbor graph. Clusters were used for downstream analyses, including visualization and trajectory inference. Clusters were annotated using known markers from described cell populations of developing testis and ovary (see also Supplemental Figures 3 and 4).

#### Trajectory inference and pseudotime analysis

Lineage relationships were inferred using partition-based graph abstraction (PAGA) implemented in Scanpy. The PAGA graph was used to guide visualization of cluster connectivity and support trajectory inference. Pseudotime was computed using diffusion pseudotime (DPT), with root cells defined based on established marker gene expression. Cells were ordered along inferred differentiation trajectories.

#### RNA velocity analysis

RNA velocity was estimated using Velocyto to quantify spliced and unspliced transcripts from Cell Ranger-generated BAM files. The resulting loom files were further processed using dynamical modeling implemented in scVelo. RNA velocity vectors were computed and projected onto the UMAP embedding to infer the directionality of cellular state transitions.

#### Visium 10x Spatial Transcriptomics

One fresh dissected adult medaka testis (21 weeks old), was embedded in OCT mounting medium, cryosectioned into a medial cross-section (6μm) and placed on a Visium Spatial Gene Expression slide from 10X Genomics. Samples were processed according to manufacturer’s instructions. Briefly, slides were stained with Hematoxylin and Eosin, and processed using the Visium Spatial Gene Expression Reagent kit (Visium HD 3’, 6.5mm). Sequencing was performed on an Illumina NextSeq 2000 with the following read configuration: 43 cycles for Read 1, 10 cycles each for the i5 and i7 indices, and 75 cycles for Read 2 with 200 million PE reads per library. Spatial transcriptomics data was processed using the Space Ranger (v4.0.1) software (10x Genomics) with default parameters. Raw sequencing data were aligned to the *Oryzias latipes* reference genome (Ensembl v107), and spatially resolved gene expression matrices were generated. Quality control, normalization, and downstream analyses, including clustering and visualization of spatial gene expression patterns, were performed using Seurat (v5.4.0) in R (v4.5.2). Default workflows and parameter settings were used for data filtering, dimensionality reduction, clustering, and spatial mapping. Spatial predictions of single-cell clusters were performed by using the RCTD approach implemented in the spacexr package (v2.2.1) and a bin size of 8μm.

#### Self-Assembling Manifold Mapping (SAMap)

To assess potential homology of cell types between medaka and other vertebrates, we compared the medaka embryonic gonad dataset with publicly available embryonic gonad datasets from mouse [64], chicken [55] and turtle [57] using the Self-Assembling Manifold Mapping algorithm (SAMap; v0.2.3) [56]. Analyses were performed in a pairwise manner (medaka–mouse and medaka–chicken).

Briefly, SAMap integrates single-cell transcriptomic atlases across species by combining sequence homology and gene expression similarity. First, a gene–gene bipartite graph was constructed by identifying homologous gene pairs between species using reciprocal BLASTP searches between their proteomes. Only reciprocal best hits were retained to define high-confidence orthologous relationships, which were used as cross-species edges in subsequent steps. Next, each dataset, independently annotated, was projected into a shared low-dimensional manifold. During this step, SAMap iteratively refines the mapping by weighting gene–gene edges based on expression correlations across aligned cells, thereby updating the homology graph and improving cross-species alignment. Finally, SAMap computes alignment scores between all pairs of cell clusters (based on species-specific annotations) across species. These mapping scores (ranging from 0 to 1) quantify the strength of correspondence between clusters based on their transcriptional similarity in the joint manifold. High-scoring pairs (>0.7) were interpreted as candidate homologous or closely related cell types.

## Acknowledgments

This research was supported by grants from the French National Research Agency (ANR-23-CE20-0025, Cell2Fish and ANR-09-GENM-017-001, AphiSwitch) to AH; (ANR-23-CE13-0008, Evo-Ovo) to F.M and “Région Bretagne” (CELL_ID) to F.M. and F.H.S.

## Author contributions

Conceptualization: AD, CM, FHS, FM, MS and AH; Data curation: AD, CM, FHS, AB, FM; Formal analysis: AD, FHS, AB, ALC, CB, FB, FM, AH; Funding acquisition: FM, AH; Investigation: AD, FM, AH; Methodology: AD, CM, FHS, AB, FM; Project administration: AD, AH; Resources: AD, ALC; Software: AD, CM, FHS, AB; Supervision: FM, SN, MS, AH; Validation: AD, CM, FHS, FB, FM, AH; Writing original draft: AD, FM, MS, AH; Review and editing: AD, CM, FHS, SN, FM, MS, AH.

## Key Resources Table

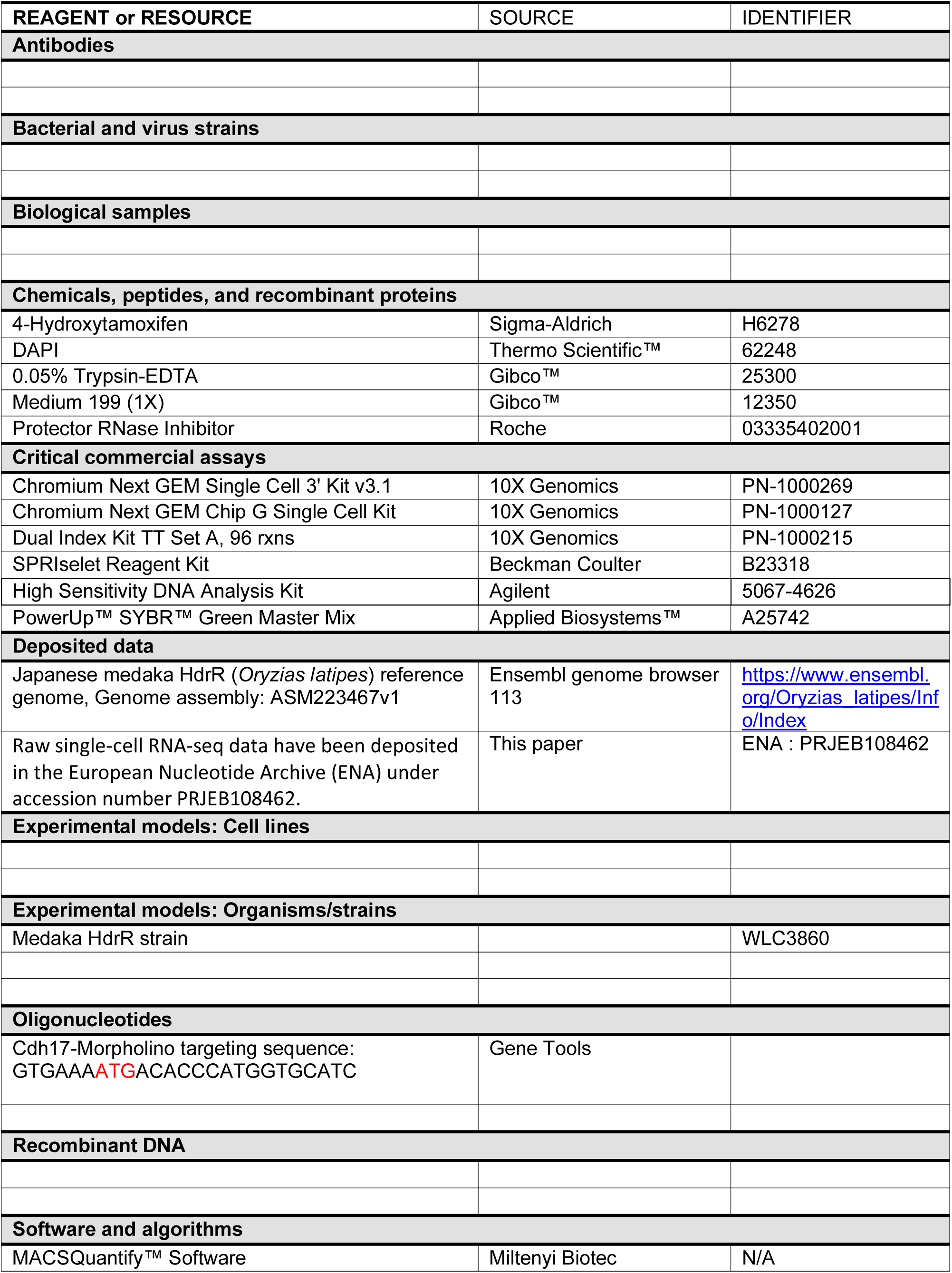
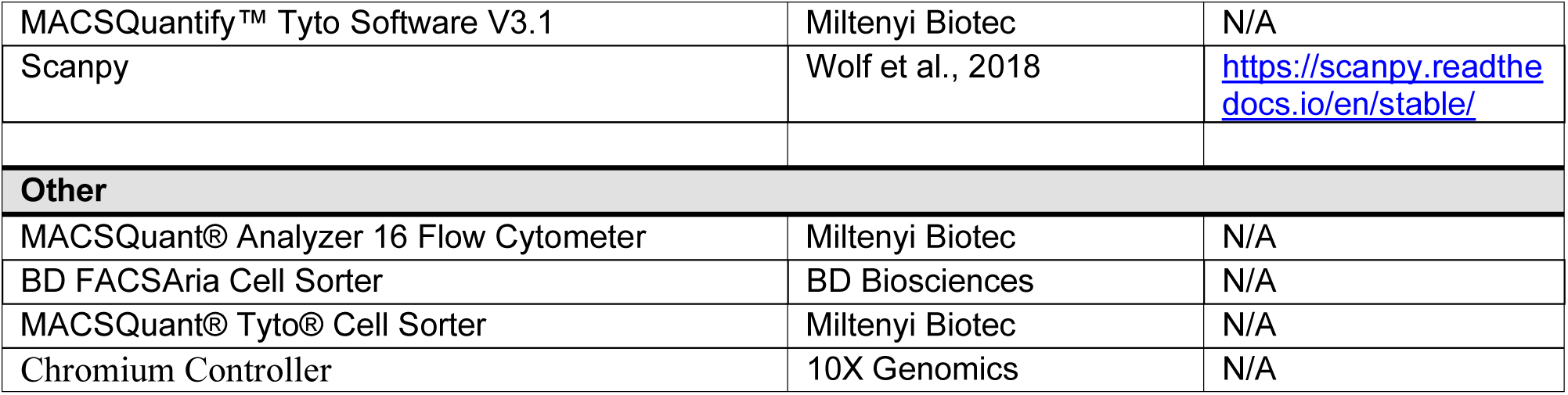

